# Plant-specific armadillo repeat kinesin directs organelle transport and microtubule convergence to promote tip growth

**DOI:** 10.1101/2022.07.08.499237

**Authors:** Asaka Kanda, Kento Otani, Takashi Ueda, Taku Takahashi, Hiroyasu Motose

## Abstract

Tip growth is essential for plant growth and reproduction. However, it remains elusive how highly polarized tip-growing cells coordinate the organization of intracellular components. Here, we show that plant-specific armadillo-repeat containing kinesin, MpARK, mediates organelle transport and microtubule convergence in tip growing rhizoids in the early diverging land plant *Marchantia polymorpha*. MpARK is required for anterograde transport of organelles to maintain their position. Furthermore, MpARK participates in the formation of microtubule foci at the rhizoid apex to stabilize growth direction. MpARK-dependent rhizoid growth is essential for plant anchorage and soil holding capacity. Thus, ARK might be a principal organelle transporter and intracellular organizer in the primitive rooting cells for the adaptation to terrestrial environments. Our findings suggest that ARK is functionally comparable to canonical anterograde kinesins in animals and fungi.

## Introduction

Tip growth is evolutionarily conserved process essential for morphogenesis and reproduction in various organisms. Furthermore, it is an excellent model system to study cell polarity and directional growth^1, 2^. During the initial stage of tip growth, one growth site is established to form a bulge in the outer surface. The bulge expands to form a dome-shaped apex followed by a cylindrical shank, resulting in a long filamentous shape. Although the detailed mechanism how a growth site is established and maintained is uncertain, local activation of ROP (Rho of plant) GTPases participates in growth site determination^3, 4^ and then actin-myosin-mediated vesicle transport and cytoplasmic streaming are thought to drive tip growth^5, 6^. However, microtubule-dependent transport have not been well clarified.

Recent studies demonstrate that kinesins, a family of microtubule-based motor proteins, play key roles in tip growth^7–13^. Several kinesins regulate microtubule organization and dynamics to promote tip growth^7, 8^. The kinesin-14 family is diverged in land plants to mediate minus end-directed transport of organelles, microtubules, actin filaments, and chromatin, most likely as the counterpart of cytoplasmic dynein in animals^10, 11, 14–17^. Two kinesin-14 subfamilies, KCBP (kinesin-like calmodulin binding protein) and KCH (kinesin with calponin homology domain), drive retrograde transport of the nucleus and plastids in protonemal apical cells in the moss *Physcomitrium patens*^10, 11^. In animals, plus-end-directed kinesins (kinesin-1, 2, 3) predominantly transport organelles toward the microtubule plus end^18–20^, whereas such kinesin remains unclear in plants.

Plant-specific armadillo-repeat kinesin (ARK) has a N-terminal motor domain followed by the C-terminal tail, which contains coiled-coil domains and tandem armadillo repeats. Because the ARK motor domain is not related to those of other kinesin families, they constitute an ungrouped orphan kinesin family^21–23^, implying their unknown evolutional origin and unpredictable function. However, microtubule gliding assay has proved the plus-end directed motor activity of *P. patens* kinesin-ARK-a^9^. Furthermore, RNAi screening has shown that *P. patens* kinesin-ARKs are required for anterograde transport of the nucleus toward the protonema apex^9^. However, ARK function remains unclear because the RNAi knockdown did not cause any obvious defect in tip growth.

Molecular genetic studies identified ARK1 of *Arabidopsis thaliana* as a regulator of root hair tip growth^24–26^. *Arabidopsis* has three *ARK* genes, *ARK1*-*ARK3*, which may redundantly and/or independently function in the cell- and organ-specific manners. The *ark1* mutant has short, branching root hairs and excess endoplasmic microtubules^25^. ARK1 localizes to the growing plus ends of microtubules to promote their catastrophe and growth velocity^7^. ARK1 interacts with NIMA (Never In Mitosis A)-related kinase 6 (NEK6)^25^ while they independently function in root hairs^27^. ARK1 interplays with the atlastin GTPase RHD3 (Root Hair Defective3) to mediate transport of ER along microtubules^28^. Previous studies have also suggested interactions of ARK1 with ROP2 and ADF1 (a class I ADP ribosylation factor GTPase-activating protein) ^29, 30^.

Rhizoid is a putative early rooting structure in land plants and is emerging as a new model of tip growth^31, 32^. A series of studies in rhizoids of *Marchantia polymorpha*, an early diverging land plant, implies common regulators in both rhizoids and root hairs^33–35^. *M. polymorpha* has several advantages including low genetic redundancy^36^ and amenable genetic tools^37, 38^. The extensive mutant screening has shown that genes involved in cell wall biogenesis, vesicle transport, and cytoskeleton are required for rhizoid growth in *M. polymorpha*^34^. We found that a tubulin kinase MpNEK1 localizes to microtubule foci at the apical dome of rhizoid to direct tip growth through microtubule reorganization^39^. Another microtubule-associated protein, MpWDL (WAVE DAMPENED-LIKE), regulates growth direction of rhizoids by stabilizing microtubules aligned along the longitudinal axis of rhizoids^40^. Here, we found that *M. polymorpha* armadillo-repeat kinesin, MpARK, directs organelle transport and microtubule convergence to promote rhizoid growth. ARK might be a predominant anterograde transporter to promote tip growth of rooting cells in land plants.

## Results

### MpARK promotes tip growth of rhizoids

A single ARK gene, Mp*ARK* (Mp4g06510), was identified in the genome database of the liverwort *M. polymorpha*. The cloned cDNA encodes a protein of 883 amino acid residues composed of a kinesin motor domain, a coiled-coil domain, and four tandem armadillo repeats, the well-conserved domain structure of plant-specific armadillo-repeat kinesin (Fig. 1a). There were seven splicing variants of MpARK, from Mp4g06510.1 to Mp4g06510.7 (Supplementary Fig. 1a). This cDNA corresponded to Mp4g06510.2, Mp4g06510.3, and Mp4g06510.7, in which the 12th and 13th exons encoding the C-terminal region of coiled-coil domain were spliced out (Fig. 1b, Supplementary Fig. 1a). The RT-qPCR analysis showed that the transcripts without the 12th and 13th exons were more abundant than the transcripts with these exons (Supplementary Fig. 1b).

**Fig. 1.**
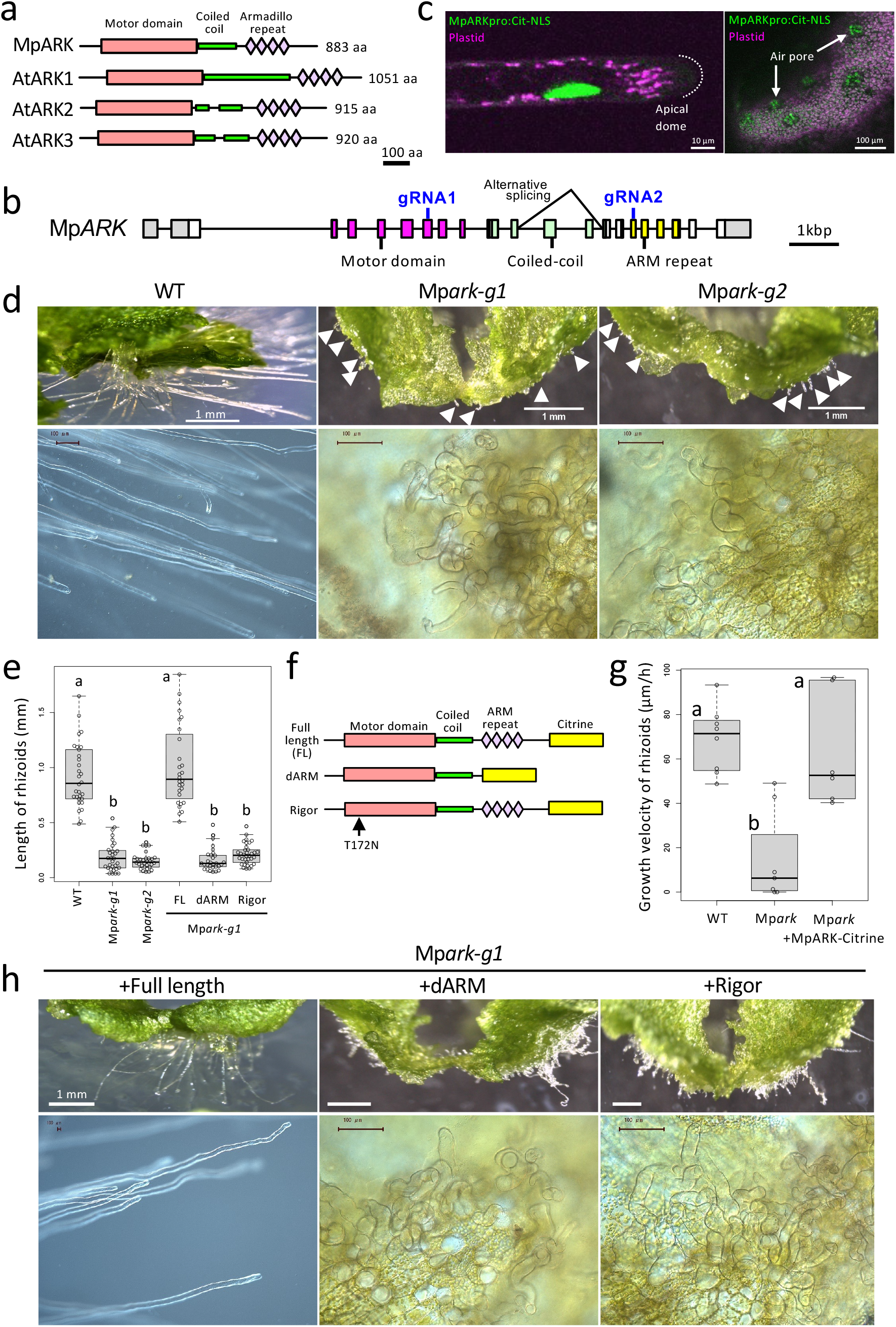
MpARK is required for tip growth of rhizoids. (a) Schematic primary structures of MpARK and AtARKs. (b) Exon-intron structure of MpARK. Box, exon; line, intron; grey, 5’ or 3’ untranslated region; magenta, motor domain; blue, coiled-coil domain; yellow, armadillo repeats. (c) The expression pattern of MpARK*pro:Citrine-NLS*. Left panel: a confocal image of Citrine-NLS (green) in the rhizoid nucleus, right panel: confocal z-stack image of Citrine-NLS (green) in the nuclei of thallus cells. Magenta, plastid autofluorescence. Arrows indicate air pore cells. (d) Morphology of 14-day-old wild type (Tak-1) and Mp*ark* mutants (Mp*ark-g1* and Mp*ark-g2*). Lower panels are images of rhizoids. Arrowheads indicate rhizoids. (e) Length of rhizoids. Values are shown by box plots indicating median (middle line), 25^th^, 75^th^ percentile (box) and 5th and 95th percentile (whiskers) (n = 28-34 rhizoids in 4-6 four-day-old plants). The different letters indicate significant differences by Tukey’s HSD test (P < 0.01). (f) Structure of MpARK constructs introduced into the Mp*ark-g1* mutant. (g) Growth velocity of rhizoids in the wild type (Tak-1), Mp*ark-g1*, and Mp*ark-g1* complemented with the full length MpARK-Citrine (MpARKpro:MpARK-Citrine). Values are shown by box plots indicating median (middle line), 25^th^, 75^th^ percentile (box) and 5th and 95th percentile (whiskers) (*n* = 6-8). The different letters indicate significant differences by Tukey’s HSD test (P < 0.01). (h) Morphology of 14-day-old Mp*ark-g1* introduced with the full length MpARK-Citrine (Full length), MpARK-Citrine without armadillo repeats (dARM), or rigor mutant MpARK-Citrine (Rigor) under the control of MpARK promoter. Lower panels show rhizoid morphology.

To analyze the expression pattern of MpARK, we generated transgenic lines expressing the nuclear-localized Citrine fluorescent protein (Citrine-NLS) under the control of the MpARK promoter (Mp*ARKpro:Citrine-NLS*). The fluorescently labeled nuclei were observed in rhizoids and in the vegetative flat organ, thallus, especially in air pore cells (Fig. 1c), implying the function of MpARK in rhizoids and thallus cells.

To address MpARK function, we generated mutants of MpARK using CRISPR/Cas9 (Fig. 1b, d, Supplementary Fig. 2, 3). The guide RNAs were designed to target a motor domain (gRNA1) and an armadillo repeat domain (gRNA2). The isolated mutants were designated Mp*ark-g1* (motor domain) and Mp*ark-g2* (armadillo repeat), respectively. All mutant lines exhibited same morphological defects in rhizoids. In the wild type, rhizoids elongated by tip growth to a filamentous shape, whereas the Mp*ark* mutants had very short, curly rhizoids (Fig. 1d, e). The phenotype is attributed to the severe suppression of tip growth and the defect of growth direction. Collectively, mutant phenotypes suggested that MpARK promotes rhizoid elongation and regulates the direction of tip growth.

In the wild type, most rhizoids entered the agar medium to fix thallus on the substratum, whereas Mp*ark* mutant rhizoids could not reach and penetrate the agar medium (Supplementary Fig. 2a). Thus, Mp*ark* mutant thallus was easily peeled away from the medium, clearly indicating that rhizoids are essential to anchor the plant body in the substratum. On the other hand, thallus growth and morphology of Mp*ark* mutants were normal compared with the wild type (Supplementary Fig. 2a). Thus, MpARK is not essential for thallus development.

### Kinesin motor and armadillo repeats are essential for MpARK function

Next, we generated transgenic lines expressing MpARK-Citrine fusion protein under the control of the MpARK promoter (Mp*ARKpro:MpARK-Citrine*) in the Mp*ark-g1* mutant background (Fig. 1 f-h). The defect of tip growth was rescued in the transgenic lines expressing MpARK-Citrine (Supplementary Fig. 2b). Rhizoid growth was equivalent between the wild type and the complementation lines (Fig. 1e, g).

We next examined the function of the kinesin motor and armadillo repeats of MpARK. The C-terminal truncated MpARK lacking armadillo repeats (dARM) and a rigor mutant without kinesin motor activity (Rigor) were tagged with Citrine and driven by the MpARK promoter in the Mp*ark-g1* mutant (Fig. 1f). The full-length MpARK tagged with Citrine recovered the defects of rhizoid growth, whereas both dARM and rigor MpARK could not (Fig. 1e, h). Collectively, both kinesin motor and armadillo repeats are essential for MpARK function in rhizoid growth.

### MpARK localizes to microtubule foci in the apical dome of rhizoids

To assess the subcellular localization of MpARK, we observed the transgenic lines expressing MpARK-Citrine by confocal microscopy. MpARK-Citrine localized to the apical dome of rhizoids, where tip growth takes place (Fig. 2a). MpARK-Citrine accumulated in the dots and along the cytoskeletal filamentous structures. MpARK dots moved toward and away from the apex with fusion and dissociation events (Supplementary Movie 1, 2, Supplementary Fig. 4a). The velocity of the apical-directed MpARK dots was 4.5±1.1 µm/min, which is almost half of that of basal-directed dots (Fig. 2b). In contrast, MpARK dARM-Citrine localized in the cytoplasm, suggesting that armadillo repeats are required for apical localization (Fig. 2c).

**Fig. 2.**
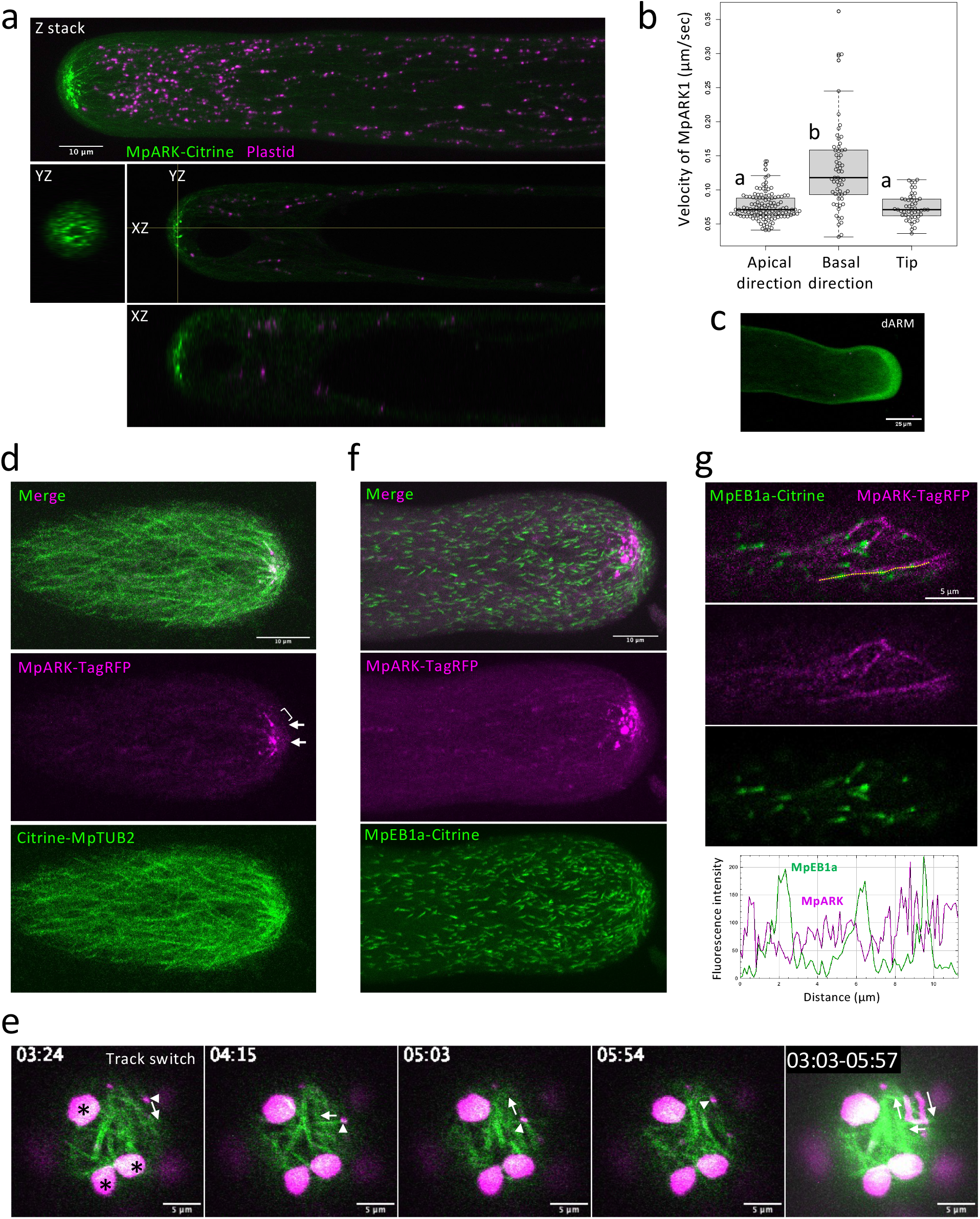
MpARK localizes in microtubule foci at the rhizoid apex. (a) Subcellular localization of MpARK-Citrine. Confocal z-stack image and orthogonal planes of a rhizoid of the complemented line (Mp*ark-g1* mutant with MpARKpro:MpARK-Citrine). (b) Velocity of MpARK-Citrine particles moving acropetally (apical direction) or basipetally (basal direction) and of MpARK-TagRFP particles along microtubules at the tip of rhizoids (tip). Values are shown by box plots indicating median (middle line), 25^th^, 75^th^ percentile (box) and 5th and 95th percentile (whiskers) [*n*=6 cells both in apical direction (*n*=119 particles) and basal direction (*n*=60 particles), *n*=5 cells in tip region (*n*=48 particles)]. The different letters indicate significant differences by Tukey’s HSD test (P < 0.01). (c) Subcellular localization of full length MpARK-Citrine and armadillo-repeats-deleted MpARK-Citrine (dARM). A single optical section of a rhizoid in Mp*ark-g1* introduced with the full length MpARK-Citrine and MpARK dARM-Citrine. (d) Localization of MpARK-TagRFP to the apical microtubules. Confocal z-stack image of double labelled line (MpARKpro:MpARK-TagRFP, CaMV35Spro:Citrine-MpTUB2). Arrows and a bracket indicate localization of MpARK-TagRFP in the microtubule ends and the side of microtubule, respectively. (e) Movement of MpARK-TagRFP particle along microtubules. MpARK-TagRFP particle switches microtubule tracks (Supplementary Movie 3). The rightmost panel, trajectory of MpARK-TagRFP. Arrowheads, MpARK-TagRFP particles. Arrows, direction of movement. Asterisks, autofluorescence of plastids (f) Different localization of MpARK-TagRFP and MpEB1a-Citrine (microtubule plus ends). Confocal z-stack image of double labelled line (MpARKpro:MpARK-TagRFP, CaMV35Spro:MpEB1a-Citrine). (g) Distinct localization of MpARK-TagRFP and MpEB1a-Citrine (microtubule plus ends). Upper panels, single cortical images in the apical region of a rhizoid of double labelled line (MpARKpro:MpARK-TagRFP, CaMV35Spro:MpEB1a-Citrine). Lower panel, fluorescence intensity profile of MpEB1a-Citrine (green) and MpARK-TagRFP (magenta) along the yellow dashed line in the uppermost panel.

We analyzed the effects of microtubule-depolymerizing drug, oryzalin, and actin filament-depolymerizing drug, latrunculin B on MpARK localization (Supplementary Fig. 4b). Treatment with oryzalin disrupted apical localization of MpARK-Citrine, resulting in dispersed fluorescence signal in the shank region. By contrast, treatment with latrunculin B did not severely affect the localization of MpARK-Citrine, still accumulating in the apical dome.

To examine microtubule localization of MpARK, we generated double labelled lines expressing both Citrine-MpTUB2 (microtubule marker, *CaMV35Spro:Citrine-*Mp*TUB2,* ref. 39) and MpARK-TagRFP (Mp*ARKpro:MpARK-TagRFP*). MpARK-TagRFP localized to microtubule foci and bundles at the apex of rhizoids (Fig. 2d, Supplementary Fig. 4c, Supplementary Movie 3). MpARK localized to the microtubule ends and laterally associated with the sides of microtubules. MpARK particles moved along microtubule lattices and switch tracks from one microtubule to another microtubule at the tip of rhizoids (Fig. 2e, Supplementary Movie 4). The particles also accumulated in the crossovers of microtubules to change their position (Supplementary Fig. 4d, Supplementary Movie 4). The velocity of MpARK particles along microtubules was equivalent to the apical-directed MpARK movement (Fig. 2b). Because most plus ends of microtubules are directed to the rhizoid apex (described below, Fig. 4f), these velocities may represent plus-end directed motility of MpARK.

Since a previous study has shown that *Arabidopsis* ARK1 localized in the microtubule plus ends^7^, we examined co-localization of MpARK-TagRFP and the plus-end marker END BINDING PROTEIN1 (*CaMV35Spro:*Mp*EB1a-Citrine*). MpARK-TagRFP did not co-localize with MpEB1a-Citrine (Fig. 2f, g, Supplementary Movie 5, 6). Consistently with this, apical-directed MpARK movement was about two-fold slower than microtubule plus-end growth visualized by MpEB1a-Citrine (described below, Fig. 4g), indicating that apical movement is not plus end tracking. Thus, MpARK is not a canonical +TIP protein preferentially accumulating in the plus ends.

To further determine precise localization of MpARK with high resolution, we observed double labelled lines of MpARK and microtubules by an enhanced-resolution confocal microscope equipped with Airyscan detector (Fig. 3a, b, Supplementary Movie 7, 8). MpARK-TagRFP localized in the apical microtubule foci, especially in the crossing overs, lateral sides, and ends of microtubules. Quantification of these localization patterns indicated that MpARK preferentially associated with the crossovers and lateral sides of microtubules, whereas microtubule end localization is relatively few.

**Fig. 3.**
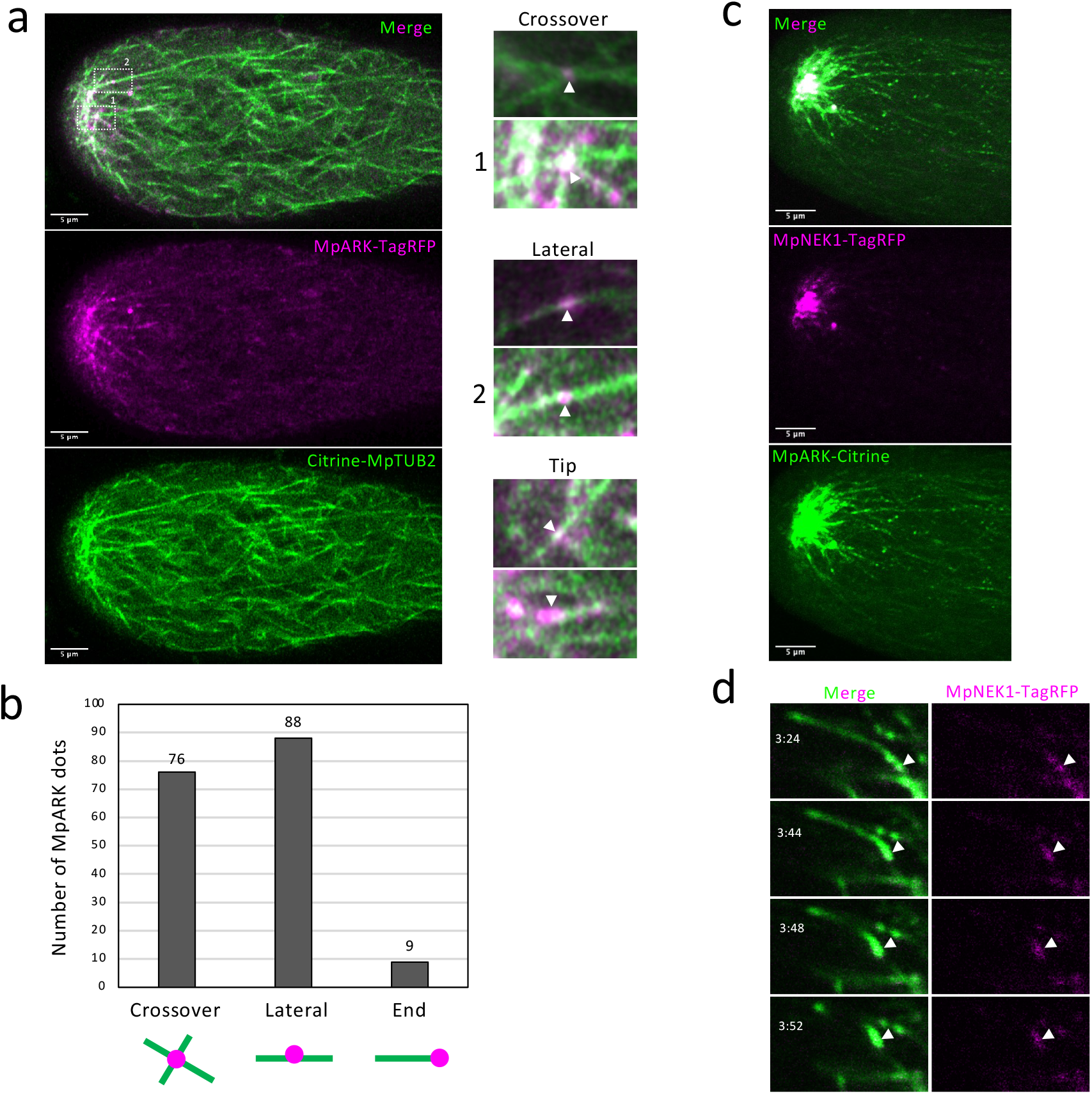
Localization pattern of MpARK in microtubules. (a) Localization of MpARK-TagRFP and microtubules. Single optical section of double labelled line (MpARKpro:MpARK-TagRFP, CaMV35Spro:Citrine-MpTUB2) observed using a confocal enhanced-resolution microscope with Airyscan detector. Right panels indicate typical localization patterns of MpARK. Arrow heads indicate MpARK-TagRFP. “1” and “2”-labelled panels are identical to the images surrounded by dotted rectangles in the left uppermost merged image. (b) Quantification of MpARK localization pattern. The numbers of MpARK dots with each localization pattern were counted in 6 rhiozids (n = 16-41 dots in each rhizoid). (c) Localization of MpARK-Citrine and MpNEK-TagRFP. Confocal z-stack image of double labelled line (MpARKpro:MpARK-Citrine, MpNEK1pro:MpNEK1-TagRFP). (d) Localization of MpARK-Citrine and MpNEK-TagRFP at a shrinking microtubule end (arrowheads).

**Fig. 4.**
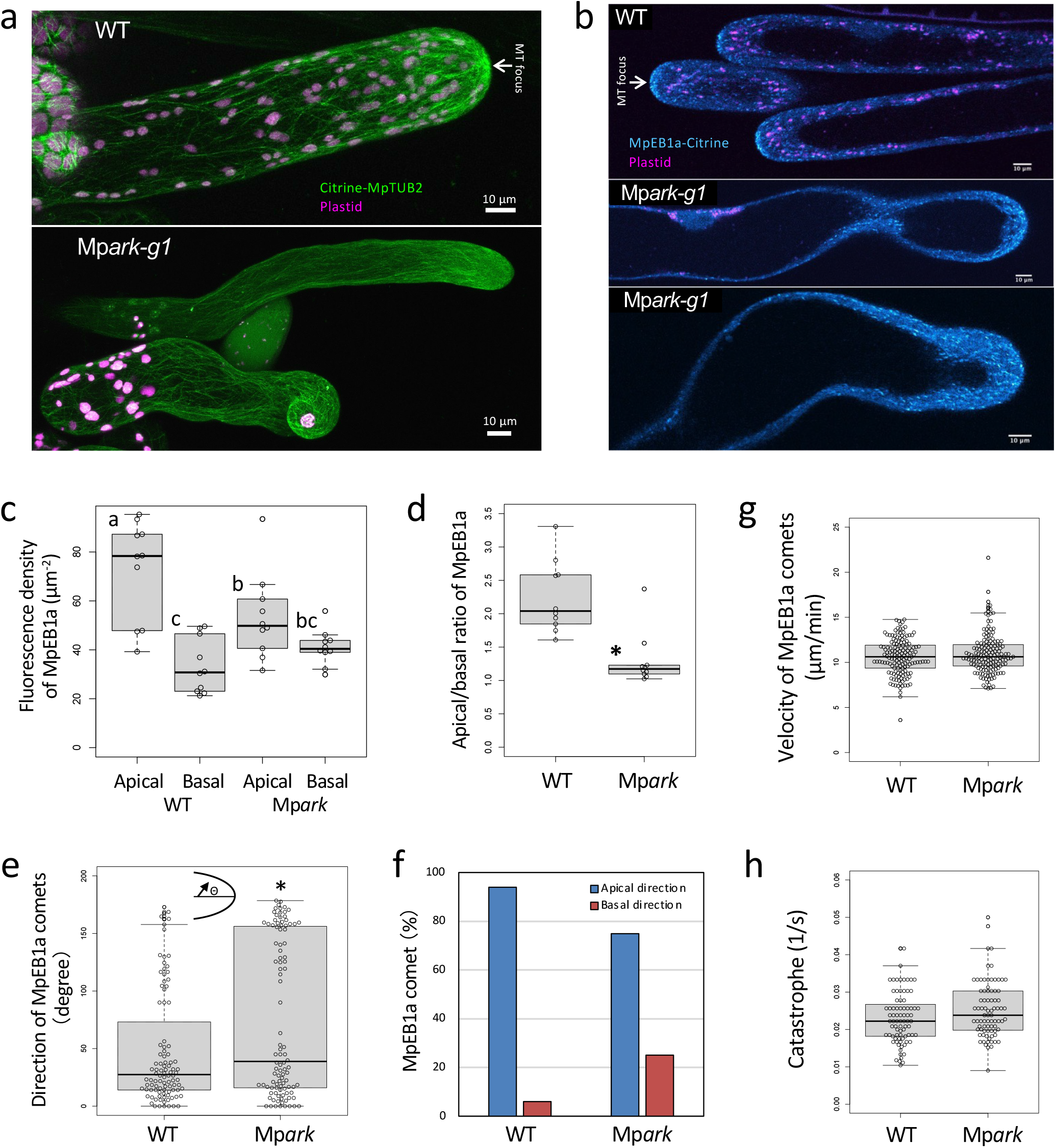
MpARK is required for microtubule convergence at the rhizoid apex. (a) Microtubules labeled with Citrine-MpTUB2 in the wild type and Mp*ark-g1*. (b) The growing plus ends of microtubules labeled with MpEB1a-Citrine in the wild type and Mp*ark-g1*. (c) Quantification of fluorescent density of MpEB1a-Citrine signal at the apical and basal regions of rhizoids in the wild type and Mp*ark-g1*. The apical region is 0-10 µm region away from the rhizoid apex and the basal region is 50-100 µm shank region away from the apex. Values are shown by box plots indicating median (middle line), 25^th^, 75^th^ percentile (box) and 5th and 95th percentile (whiskers) (*n*=10 cells). The different letters indicate significant differences by Tukey’s HSD test (P < 0.04). (d) Ratio of MpEB1a-Citrine signal at the apical and basal regions of rhizoids in the wild type and Mp*ark-g1*. Values are shown by box plots indicating median (middle line), 25^th^, 75^th^ percentile (box) and 5th and 95th percentile (whiskers) (*n*=10 cells). An asterisk indicates significant difference from the value of the wild type (Wilcoxon rank sum test, P < 0.001). (e) Direction of movement of MpEB1a-Citrine comets. The angle indicates a deviation from the longitudinal axis of rhizoids. Values are shown by box plots indicating median (middle line), 25^th^, 75^th^ percentile (box) and 5th and 95th percentile (whiskers) (*n*=100 comets, 5 cells). An asterisk indicates significant difference from the value of the wild type (Wilcoxon rank sum test, P < 0.03). (f) Proportions of MpEB1a-Citrine comets moving toward the apex or the base (*n*=100 comets, 5 cells). (g) Velocity of MpEB1a-Citrine comets. Values are shown by box plots indicating median (middle line), 25^th^, 75^th^ percentile (box) and 5th and 95th percentile (whiskers) (wild type *n*=151, 10 cells, Mp*ark n*=155, 8 cells, Wilcoxon rank sum test, P > 0.05). (h) Catastrophe of microtubule plus ends. The catastrophe of the plus ends was evaluated by the reciprocal of MpEB1a-Citrine comet duration time (time from appearance to disappearance of EB1 comet). Values are shown by box plots indicating median (middle line), 25^th^, 75^th^ percentile (box) and 5th and 95th percentile (whiskers) (*n*=80, 5 cells, Wilcoxon rank sum test, P > 0.05).

We previously showed that a NIMA-related kinase, MpNEK1, localizes to the apical microtubule foci to regulate growth direction of rhizoids through microtubule reorganization^39^. To further characterize the localization of MpARK, we generated the double labelled lines expressing both MpARK-Citrine and MpNEK1-TagRFP (Fig. 3c, d, Supplementary Movie 9, 10). MpARK colocalized with MpNEK1 in microtubule foci at the apex of rhizoids. Both MpARK and MpNEK1 associated with microtubule bundles, crossing overs, and shrinking ends. Collectively, our results suggest that MpARK localizes to microtubule foci, crossovers, lateral sides, and shrinking ends to regulate microtubule organization during tip growth of rhizoids.

### MpARK is required for microtubule convergence in the rhizoid apex

Because *Arabidopsis* ARK1 regulates microtubules in root hairs^7, 25^, we assumed that MpARK also regulates microtubule organization and dynamics. To visualize microtubules in rhizoids, we introduced a microtubule marker Citrine-MpTUB2 or a growing plus-end marker MpEB1a-Citrine in the wild type and Mp*ark-g1* mutants (Fig. 4).

In the wild type, microtubules were parallel or oblique to the longitudinal axis of rhizoids and converged into the focus at the rhizoid apex (Fig. 4a). On the while, the Mp*ark* mutants did not form obvious microtubule foci at the apex.

MpEB1a-Citrine moved toward the rhizoid apex to converge on the focus in the wild-type, whereas it did not coalesce at the apex in the Mp*ark* mutant (Fig 4b, Supplementary Movie 11, 12). The fluorescent density of MpEB1a-Citrine was decreased in the apical dome of Mp*ark* rhizoids (Fig. 4c, d). We assessed the direction of MpEB1a-Citrine comets and found its significant dispersion in Mp*ark* mutants (Fig. 4e). In the Mp*ark* mutant, basal-directed plus ends were increased but apical-directed plus ends were still dominant (Fig. 4f). The velocity of MpEB1a-Citrine comets and catastrophe of plus ends (frequency of growth-to-shrink transition evens) was not altered in the Mp*ark* mutant (Fig. 4g, h). These results indicate that MpARK regulates microtubule convergence at the rhizoid apex.

We next examined whether MpARK regulates apical localization of MpNEK1, a regulator of apical microtubule foci during tip growth of rhizoids^39^ (Supplementary Fig. S5). MpNEK1-Citrine localized to microtubule foci in the wild type but dispersed to the shank region in Mp*ark-g1*, suggesting that MpARK regulates microtubule convergence through the transport and localization of MpNEK1.

### MpARK is required for anterograde transport of organelles

We found abnormal localization of plastids and the nucleus in MpARK rhizoids (Fig. 4a, 4b, 5a). A previous study demonstrates that *P. patens* ARK kinesins are required for the movement of the nucleus to the apical direction of protonema after mitosis^9^. Thus, we analyzed the movement of the nucleus and plastids during rhizoid growth. In the wild type, the nucleus moved toward the rhizoid apex at the same velocity of rhizoid growth and kept the distance about 60-100 µm apart from the apex (Fig. 5a-c, Supplementary Movie 13). In the Mp*ark* mutants, nuclei were left behind in the basal region of rhizoids due to the significant decrease of anterograde movement (Fig. 5a-c, Supplementary Movie 14).

**Fig. 5.**
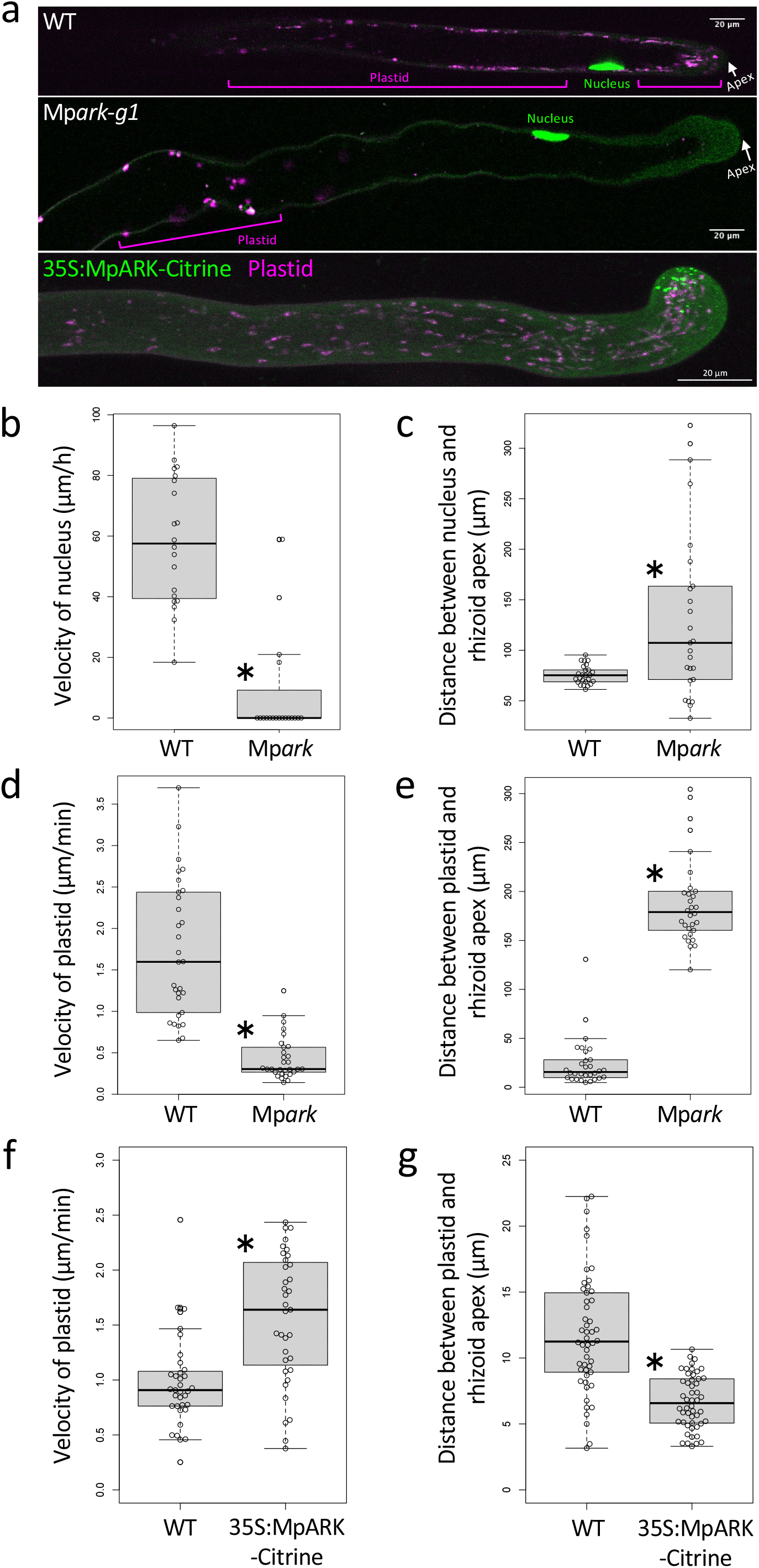
MpARK is required for organelle transport in rhizoids. (a) Localization of the nucleus and plastids in rhizoids of the wild type and Mp*ark-g1* (upper and middle panels). Localization of plastids and MpARK-Citrine in CaMV35S:MpARK-Citrine (lower panel). (b, c) Velocity of nuclear movement (b) and distance between the nucleus and the rhizoid apex (c) in the wild type and Mp*ark-g1*. The movement of nuclei labelled with Citrine-NLS (MpNEK1pro:Citrine-NLS) was observed in a confocal microscope. Values are shown by box plots indicating median (middle line), 25^th^, 75^th^ percentile (box) and 5th and 95th percentile (whiskers) (*n* = 20 nuclei in (b), *n* = 25 nuclei in (c)). An asterisk indicates significant difference from the value of the wild type (Wilcoxon rank sum test, P < 0.01). (d, e) Velocity of plastid movement (d) and distance between plastids and the rhizoid apex (e) in the wild type and Mp*ark-g1*. Five plastids were analyzed in each cell (*n* = 6 cells). The five plastids nearest to the apex were measured in (e). Values are shown by box plots indicating median (middle line), 25^th^, 75^th^ percentile (box) and 5th and 95th percentile (whiskers) (*n* = 30 plastids). An asterisk indicates significant difference from the value of the wild type (Wilcoxon rank sum test, P < 0.01). (f, g) Velocity of plastid movement (f) and distance between plastids and the rhizoid apex (g) in the wild type and CaMV35S:MpARK-Citrine. Four to eight plastids were analyzed in each cell in (f) (*n* = 35 plastids in 6 cells). The five plastids nearest to the apex were measured in each cell in (g) (*n* = 50 plastids in 10 cells). Values are shown by box plots indicating median (middle line), 25^th^, 75^th^ percentile (box) and 5th and 95th percentile (whiskers) (*n* = 35 in (f) and 50 in (g)). An asterisk indicates significant difference from the value of the wild type (Wilcoxon rank sum test, P < 0.01).

We next analyzed the movement of plastids. In the wild type, plastids showed the back-and-forth movement, but proceeded toward the apex to distribute uniformly from the shank to the apical region (Fig. 4a, 4b, 5a, Supplementary Movie 13). In the Mp*ark* mutants, the velocity of plastid movement was severely decreased (Fig. 5d) and most plastids were localized in the basal region behind the nucleus (Fig. 4a, 4b, 5a, 5e, Supplementary Movie 14).

If MpARK transports plastids, overexpression of MpARK may promote tip-directed movement and apical localization of plastids. To assess this possibility, MpARK-Citrine fusion protein was expressed under the control of Cauliflower mosaic virus 35S (CaMV35S) promoter in the wild type background. The velocity of plastids was significantly increased in the rhizoids of 35S:MpARK-Citrine transgenic plants compared with the wild type (Fig. 5f). Furthermore, plastids were more apically localized in the rhizoids of 35S:MpARK-Citrine (Fig. 5g). We found aberrant morphology of rhizoids in the 35S:MpARK-Citrine transgenic lines (n = 8/21 wavy, crooked rhizoids, Fig. 5a, Supplementary Fig. 6)

It was difficult to detect the MpARK-Citrine signal associated with the nucleus and plastids because organelle-associated signal was relatively weak compared with apical MpARK-Citrine signal and plastid autofluorescence was too high. However, by observing complementation lines with high expression of MpARK-Citrine, we could observe the filamentous signal of MpARK-Citrine associated with the plastids (Supplementary Fig. 7). To separate the MpARK-Citrine signal and plastid autofluorescence, we conducted spectral imaging combined with linear unmixing (Supplementary Fig. 8). Although cytoplasmic MpARK-Citrine signal was high, filamentous signals of MpARK-Citrine were associated with plastids. On the other hand, it was more difficult to observe the MpARK-Citrine signal associated with the nucleus. We merely detected cytoplasmic and thin filamentous signals of MpARK-Citrine around the nucleus (Supplementary Fig. 7).

To determine whether MpARK regulates movement of other organelles, we investigated *trans*-Golgi network (TGN). TGN was labelled with Citrine-MpSYP6A^41^ in the wild type and Mp*ark-g1* and observed under a confocal microscope (Fig. 6a). MpSYP6A-labelled TGN exhibited back-and-forth movement and moved to the rhizoid apex in the wild type (Fig. 6, Supplementary Movie 15). In the Mp*ark-g1* rhizoids, motility of TGN was severely reduced and most of them showed Brownian motion (Fig. 6b, Supplementary Movie 16).

**Fig. 6.**
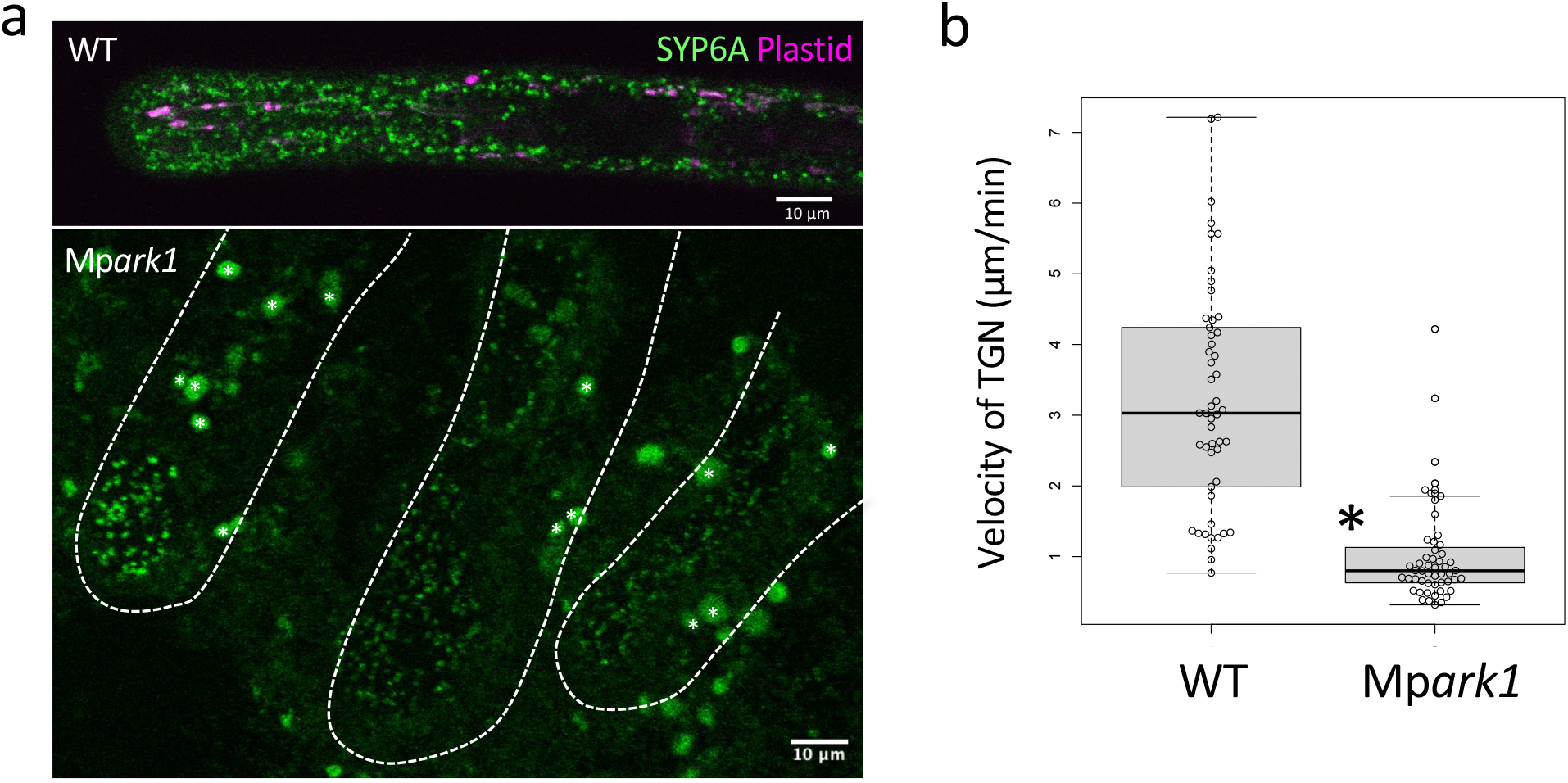
MpARK is required for TGN transport in rhizoids. (a) Localization of TGN labelled with Citrine-MpSYP6A in rhizoids of the wild type and Mp*ark-g1*. Asterisks indicate plastid autofluorescence. (b) Velocity of TGN in the wild type and Mp*ark-g1*. The movement of TGN labelled with Citrine-MpSYP6A was observed in a confocal microscope. Values are shown by box plots indicating median (middle line), 25^th^, 75^th^ percentile (box) and 5th and 95th percentile (whiskers) (*n* = 50 TGN in 3 cells in the wild type and 52 TGN in 5 cells in Mp*ark-g1*). An asterisk indicates significant difference from the value of the wild type (Wilcoxon rank sum test, P < 0.01).

We further analyzed localization of mitochondria and vesicles by staining with DiOC6 and BODIPY-conjugated brefeldin A (BODIPY-BFA), respectively (Supplementary Fig. 9). Interestingly, DiOC6-positive mitochondria and BODIPY-BFA-positive vesicles were preferentially accumulated at the apical dome in the wild type rhizoids, whereas they were dispersed in the Mp*ark-g1* rhizoids (Supplementary Fig. 9a, c). Their apical signal was decreased in Mp*ark-g1* compared with the wild type (Supplementary Fig. 9b, d). Thus, MpARK may drive anterograde transport of various organelles to coordinate their subcellular distribution during tip growth of rhizoids.

### MpARK-mediated rhizoid growth is required for holding the soil and promoting vegetative/reproductive development

To address functional significance of MpARK-mediated rhizoid growth, we monitored plant growth on soil. On the agar medium, the vegetative flat organ, thallus, mainly constituting the liverwort body, grew normally in the Mp*ark* mutant compared with the wild type (Supplementary Fig. 2a). This is likely due to direct absorption of nutrients and water from thalli and enough humidity in the closed culture plates. These artificial environments may mask the requirement of rhizoids and MpARK function.

We planted the wild type, Mp*ark* mutants, and Mp*ark-g1* mutants introduced with full length, dARM, and rigor MpARK tagged with Citrine on the soil (Fig. 7a). Thallus growth was suppressed in Mp*ark* mutants, whereas it was recovered in the Mp*ark-g1* expressing full length MpARK-Citrine. The growth suppression could not be rescued in Mp*ark-g1* introduced with dARM or rigor MpARK.

**Fig. 7.**
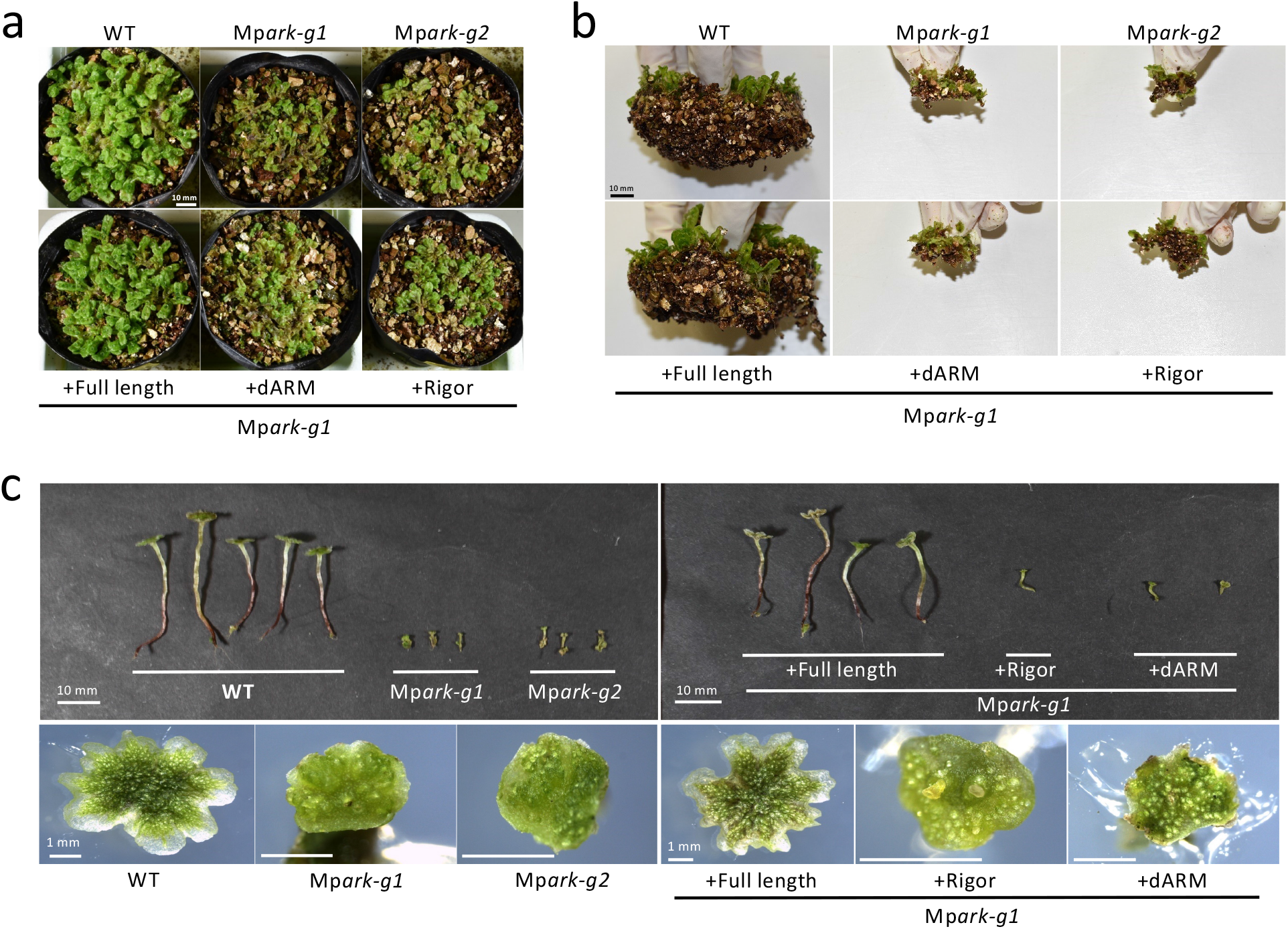
MpARK-mediated rhizoid growth is required for soil holding and reproductive development. (a, b) Thallus growth (a) and soil-holding capacity (b) of the wild type, Mp*ark-g1*, Mp*ark-g2*, and Mp*ark-g1* with the full length MpARK-Citrine, armadillo-repeats-deleted MpARK-Citrine (dARM), or rigor mutant MpARK-Citrine (Rigor). Plants were grown on the soil for one month. (c) Morphology of male reproductive organs (antheridiophores) of the wild type, Mp*ark-g1*, Mp*ark-g2*, and Mp*ark-g1* with the full length MpARK-Citrine (Full length), armadillo-repeats-deleted MpARK-Citrine (dARM), or rigor mutant MpARK-Citrine (Rigor).

We next monitored soil-holding capacity by lifting thalli up in the air (Fig. 7b). Comparing with the wild type and Mp*ark-g1* complemented with full length MpARK, Mp*ark-g1*, Mp*ark-g2* and Mp*ark-g1* introduced with dARM or rigor MpARK exhibited severely reduced soil-holding capacity, implying that the growth suppression is attributed to the reduced capacity holding the soil and the resulting defect in absorption of nutrients and water.

We next examined reproductive development (Fig. 7c). The wild type and Mp*ark-g1* complemented with MpARK developed fully grown reproductive organ (antheridiophore, male reproductive organ), whereas Mp*ark* mutants and Mp*ark-g1* introduced with dARM or rigor MpARK showed severe dwarf phenotype of antheridiophores. Thus, MpARK is required for reproductive development.

### Evolution of *ARK*

Phylogenetic analysis showed that land plant ARKs can be classified into two clades: the AtARK2 (ARK2) and AtARK1 (ARK1) clade (Fig. 8a), consistently with previous studies^21, 23^. MpARK belongs to the AtARK2 clade like as do ARKs of the moss *P. patens*, lycophyte, monilophytes, and gymnosperms, implying that the *ARK2* clade was ancestral.

**Fig. 8.**
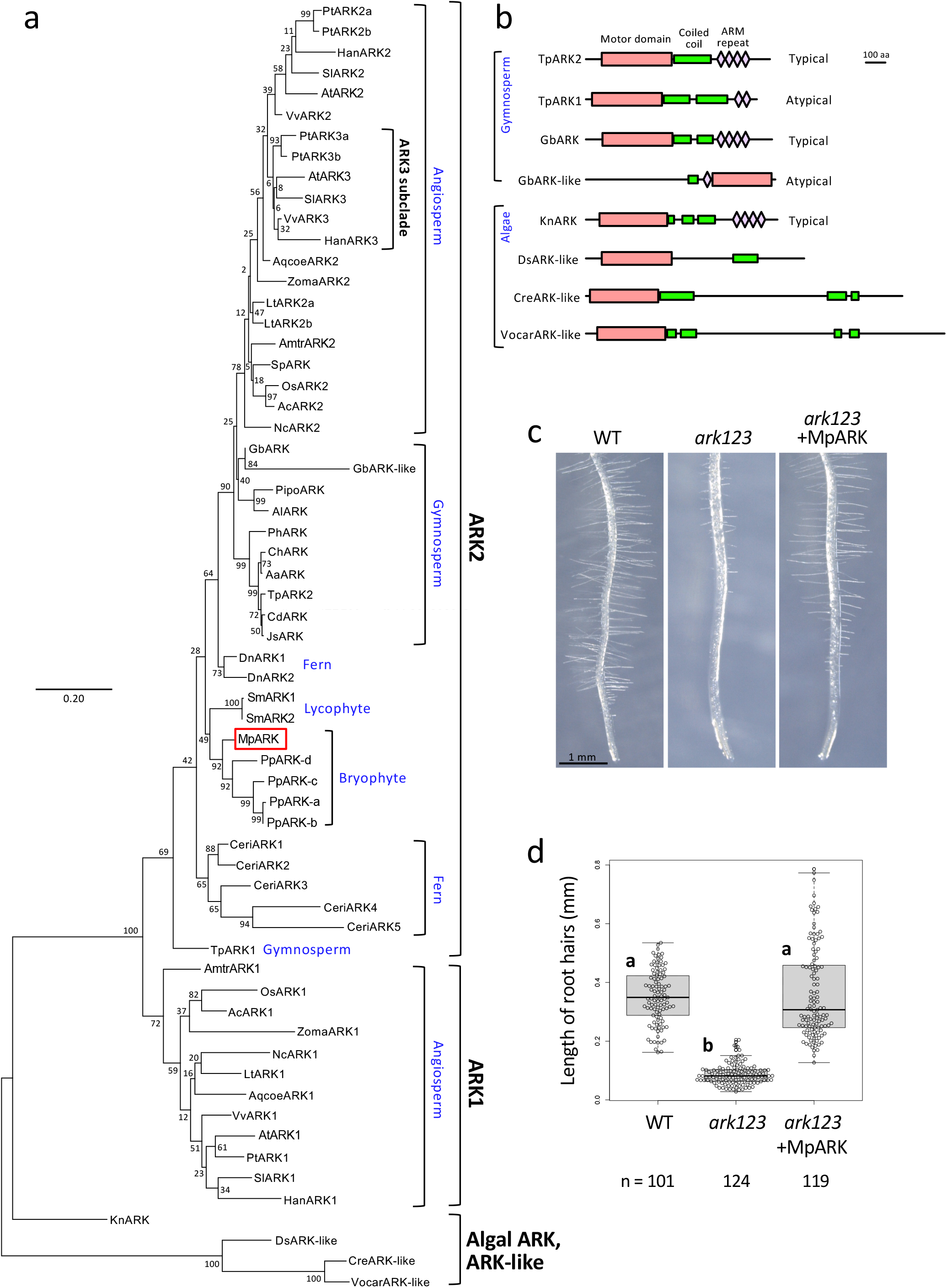
Phylogenetic analysis and functional conservation of ARK. (a) A maximum-likelihood tree constructed using the kinesin motor domain and the flanking region (corresponds to the amino acid residues 40-494 of MpARK). Numbers on the branch nodes show bootstrap support values calculated from 1000 replicates. Aa: *Amentotaxus argotaenia*, Ac: *Ananas comosus*, Al: *Abies lasiocarpa*, Amtr: *Amborella trichopoda*, Aqcoe: *Aquilegia coerulea*, At: *Arabidopsis thaliana*, Cd: *Calocedrus decurrens*, Ceri: *Ceratopteris richardii*, Ch: *Cephalotaxus harringtonia*, Cre: *Chlamydomonas reinhardtii*, Dn: *Danaea nodosa,* Ds: *Dunaliella salina*, Gb: *Ginkgo biloba*, Han: *Helianthus annuus*, Js: *Juniperus scopulorum*, Kn: *Klebsormidium nitens*, Lt: *Liriodendron tulipifera*, Mp: *Marchantia polymorpha*, Nc: *Nymphaea colorata*, Os: *Oryza sativa*, Ph: *Phyllocladus hypohyllus*, Pipo: *Pinus ponderosa*, Pp: *Physcomitrium patens*, Pt: *Populus trichocarpa*, Sl: *Solanum lycopersicum,* Sm: *Selaginella moellendorffii,* Sp: *Spirodela polyrhiza,* Tp: *Thuja plicata,* Vocar: *Volvox carteri,* Vv: *Vitis vinifera,* Zoma: *Zostera marina*. (b) Protein structures of ARKs of gymnosperms and algae. (c) MpARK rescues the growth defect of root hairs in *Arabidopsis ark1 ark2 ark3* mutant. The roots of the wild type and *ark1-1 ark2-1 ark3-1* mutant (*ark123*) with or without At*NEK6pro:*MpARK*-Citrine* were observed in a stereomicroscope. (d) The lengths of root hairs of the wild type and *ark1-1 ark2-1 ark3-1* mutant (*ark123*) with or without At*NEK6pro:*MpARK*-Citrine*. Values are shown by box plots indicating median (middle line), 25^th^, 75^th^ percentile (box) and 5th and 95th percentile (whiskers). Five plants in each genotype were analyzed. n, number of root hairs. The different letters indicate significant differences by Tukey’s HSD test (P < 0.001).

*ARK1* genes were found in most angiosperms but not in bryophytes, lycophyte, monilophytes, gymnosperms (Fig. 8a). The early diverging angiosperm *Amborella trichopoda* have two *ARK* genes, each of which belongs to the *AtARK2* or *AtARK1* clade. An exception in angiosperms is a duckweed *Spirodela polyrhiza*, which has an *ARK2*-clade gene but no *ARK1* gene. It is noteworthy that *S. polyrhiza* lacks root hairs and genes required for root hair development^42^. *ARK3* constitutes a subclade of *ARK2* (*ARK3* subclade). *ARK3* genes were specifically found in core eudicots. Thus, we conclude that *ARK1* emerged in angiosperms whereas *ARK3* evolved in core eudicots probably from the duplicated *ARK2*.

We found a typical ARK with a N-terminal motor domain and tandem armadillo repeats in a charophyte alga *Klebsormidium nitens,* whereas non-charophyte algae have no canonical ARK (Fig. 8a, b). However, we found kinesin genes with an ARK-like motor domain but without obvious armadillo repeats in non-charophyte algae. These genes may represent the early diverged ARK-like kinesin or may evolved uniquely in these algae.

Interestingly, we also noticed unusual *ARK*-like genes in gymnosperms (Fig. 8b). *Thuja plicata* ARK1 (TpARK1) has a N-terminal ARK motor domain and two armadillo repeats. *Ginkgo biloba* ARK-like (GbARK-like) has an armadillo-like domain and a C-terminal ARK-like motor domain. From the phylogenetic relationships, GbARK-like may be recently occurred from a duplicated gene, whereas TpARK1 seems to be early diverged.

To examine the functional conservation of ARK in tip growth, we introduced MpARK in *Arabidopsis ark1 ark2 ark3* triple mutant, which exhibits short, branched root hairs. The defect of root-hair tip growth was rescued by MpARK (Fig. 8c, d), suggesting the conserved ARK function in tip growth of rooting cells in land plants.

## Discussion

Here, we show that the early diverging land plant *M. polymorpha* has a single ARK kinesin (MpARK), which is essential to promote tip growth of rhizoids, the ancestral rooting system in land plants. Our study provides novel insights on the fundamental function of this plant-unique kinesin because most land plants, including mosses, lycophytes, and angiosperms, have multiple *ARK* genes, which redundantly work in plant development and microtubule-dependent processes.

Mp*ark* mutants showed severe growth defects in rhizoids: suppression of elongation and fluctuation of growth direction. Thus, MpARK has the dual function to promote rhizoid elongation and stabilize growth polarity. This is reminiscent of the function of *Arabidopsis* ARK1 in root hairs^7, 25^. ARK kinesin may play the evolutionarily conserved roles in tip growth of land plants. The recovery of *Arabidopsis ark* mutant by MpARK strongly supports this notion. Although the origin of this plant-specific kinesin is not clear, ARK kinesin may be helpful to promote the development of rooting cells for the colonization of terrestrial environments.

How does MpARK regulate these processes? Our functional analysis strongly suggest that MpARK plays a pivotal role in organelle transport to enhance tip growth of rhizoids. In the absence of MpARK, anterograde movement of the nucleus, plastids, and TGN are significantly decreased, and localization of various organelles are shifted to the basal side of rhizoids, which may in turn severely affects cellular activity required for tip growth (e.g., location of molecules such as RNAs, proteins, and metabolites, and targeted secretion of membranes and cell wall components to the apex). The association of MpARK-Citrine with plastids is consistent with this. Furthermore, Yosida et al.^43^ also demonstrate that *P. patens* ARK kinesins drive organelle transport to promote tip growth of protonemata. Therefore, ARK may represent anterograde transporter of organelles to sustain tip growth in land plants.

We also found that MpARK participates in microtubule convergence at the tip to stabilize growth direction. Mp*ark* mutant rhizoids exhibit no obvious microtubule foci and dispersed growth direction of microtubule plus ends. Since MpARK-Citrine preferentially associates with microtubule foci, crossovers, and lateral sides, MpARK may bundle microtubules at the apex like as another plant specific kinesins KINIDa/b in *P. patens* protonemata^6^. Because MpARK colocalizes with microtubule depolymerzing protein MpNEK1 and exhibits dynamic behavior on microtubules, MpARK may also arrange connections among microtubules to reorganize their network during tip growth. Alternatively, but not exclusively, MpARK may transport MpNEK1 to regulate the apical microtubule foci.

The roles of MpARK in organelle transport and microtubule organization might be mutually independent. Because of spatial separation between organelle transport (mainly at the shank region) and microtubule foci (at the apex), it is unlikely that MpARK regulates organelle transport through microtubule configuration. Although most microtubule plus-ends still direct to the rhizoid apex in the Mp*ark* mutant, we could not exclude that minor microtubules with aberrant directions were predominantly used as the tracks to affect organelle transport. Another regulator of rhizoids, MpNEK1, localizes to microtubule foci and stabilizes growth direction by depolymerizing and reorganizing apical microtubules^39^. Mp*nek1* mutants show fluctuation of growth direction and defect of apical microtubules, but do not exhibit abnormal organelle localization. Thus, apical microtubule foci may not be required for organelle transport but rather stabilize direction of tip growth.

Previous studies have demonstrated that *Arabidopsis* ARK1 localizes to the growing plus-ends and promotes microtubule catastrophe during root hair elongation^7^. ARK1-mediated microtubule catastrophe is presumed to increase cytoplasmic free tubulin and growth velocity of microtubule plus ends. Our study indicates no obvious defects of Mp*ark* mutants in microtubule dynamics. Thus, we inferred that the severe defects in rhizoid growth could not be attributed to microtubule dynamics. *Arabidopsis* ARK1 may acquire a novel function to regulate microtubule dynamics for optimizing root hair growth, as suggested from the divergence of ARK1 clade in angiosperms. The functional conservation and diversification among ARK subfamilies are open question for future research.

Recent studies demonstrate that kinesin-14 drives minus end-directed transport of organelles and cytoskeletons^10, 11, 14–17^. KCH drives retrograde transport of organelles in tip growing protonemata in *P. patens*^11^, whereas ARK kinesins drive anterograde transport of the nucleus^9^ and various organelles^43^. Although it remains unknown whether kinesin-14 functions in rhizoids, ARK and kinesin-14 may competitively function in organelle transport to coordinate intracellular organization. Because cell growth causes extensive displacement and rearrangement of organelles, kinesin-mediated organelle relocation might be crucial to arrange intracellular organization for stable cellular activity and growth.

Collectively, our results suggest that ARK corresponds to the canonical anterograde kinesins in animals which execute long-range transport of organelles^18–20^. ARK might be co-opted as a long-range transporter to enhance the persistent tip growth of rooting cells during land plant evolution. This coincides with the elongation of ER tubules along microtubules^44^ and ARK1-mediated ER movement to the microtubule plus ends^27^ in *Arabidopsis*. In *Arabidopsis*, myosin XI-i promotes nuclear motility in the mature cells through interaction with nuclear membrane proteins WIT1/2^45^, whereas rapid acropetal nuclear movement in growing root hairs could occur in the absence of myosin XI-i and microtubules are required for stable nuclear migration^46^, suggesting that both actin- and microtubule-based motor proteins participate in this process. Although myosin function remains unclear in *M. polymorpha*, molecular activity of myosin XI is well conserved, but the velocity of cytoplasmic streaming is very low compared with that in flowering plants^47^. Hence, actin-myosin system is not the main transport machinery in rhizoids. Because tip localization of MpARK was not affected by actin depolymerization, actin-myosin may not participate in MpARK localization and transport. However, we could not fully exclude the possibility that MpARK participates in actin-microtubule crosstalk as shown in KCH^16, 48–49^, KCBP^50^, and myosin VIII^51^. Further studies of plant kinesins and their cargos/interactors will reveal the microtubule-based transport system and its functional significance.

## Methods

### Plant Material and Growth Conditions

*M. polymorpha* accessions, Takaragaike-1 (Tak-1, male) and Tak-2 (female), were used as the wild-type. Thalli and gemmalings were grown at 22°C under continuous white light on the half-strength Gamborg’s B5 medium [Gamborg’s B5 medium mixed salt (Fujifilm Wako Pure chemical, Osaka, Japan), 0.5 g/L 2-morpholinoethanesulfonic acid (MES), 0.1 g/L myo-inositol, 0.001 g/L nicotinic acid, 0.001 g/L pyridoxine hydrochloride, 0.01 g/L thiamine hydrochloride, adjusted to pH 5.5 with KOH and solidified with 1% agar]. Antibiotics, claforan (100 mg/L), hygromycin (10 mg/L), and chlorsulfuron (5 µM) were added to the half B5 medium. For pharmacological analysis, oryzalin and latrunculin B were dissolved in dimethyl sulfoxide (DMSO) and added to B5 media as final concentrations of DMSO were less than 0.2%. Dormant gemmae were collected and plated on B5 media with or without drugs. For the estimation of plant growth, gemmalings grown for one week were transferred on the expanded vermiculite (NITTAI Co., Osaka, Japan) in the plastic pots (89 mm diameter, 75 mm height) and photographed after one month by a single-lens reflex camera D5600 (Nikon, Tokyo, Japan). The soil-holding capacity was estimated by lifting thalli (grown for one month on soil) up in the air as described previously^40^. Gametangiophore was induced under continuous white light with far-red irradiation as described previously^52^. *A. thaliana* Columbia accession was used as the wild type. The *ark1-1 ark2-1 ark3-1* triple mutant of *A. thaliana* were described previously^25^. Seeds of *A. thaliana* were germinated on the half-strength Murashige and Skoog (MS) agar medium and vertically incubated at 22°C under continuous white light as described previously^53^.

### Plasmid construction and transformation

For the mutagenesis of MpARK, we generated CRISPR/Cas9 constructs. Two complementary DNA oligos encoding 20-base target sequences of gRNA (Table S1) were annealed and cloned into the *Bsa*I site of an entry vector pMpEn_03 according to ref. 49. The Mp*U6-1pro:gRNA* sequences in pMpEn_03 plasmids were transferred into a binary vector pMpGE010^54^ by LR reaction using Gateway LR Clonase II enzyme mix (Thermo Fisher Scientific, Tokyo, Japan). The resulting binary vectors were introduced into *Agrobacterium* GV3101 (pMP90) strain by electroporation. The constructs were introduced into wild type F1 sporelings derived from sexual crosses between Tak-2 and Tak-1 by *Agrobacterium*-mediated transformation method^37^. Transformants were selected on the half B5 agar medium containing 10 µg/ml hygromycin B and 100 µg/ml claforan. For the selection of mutants, genomic DNA of transformants was extracted and subjected to PCR by a DNA polymerase KOD FX neo (Toyobo) and gene specific primers to amplify DNA regions flanking the target sequences. The primers used for PCR are shown in Table S1. By using amplified DNA as a template, sequence reaction was performed by BigDye terminator ver.3.1 (Thermo Fisher Scientific) and a gene-specific primer (Table S1). DNA sequencing was performed by ABI3500 Genetic Analyzer (Thermo Fisher Scientific).

To construct MpARKpro:MpARK-Citrine, the full-length MpARK was amplified from Tak-1 cDNA by PCR using KOD plus (TOYOBO) with the primers in Table S1 and cloned into pENTR/D-TOPO cloning vector (Thermo Fisher Scientific). This vector was confirmed by sequencing and designated as pENTR/D-TOPO-MpARK. The MpARK promoter region, including a 3.4-kb region upstream of the initiation codon, was amplified from Tak-1 genomic DNA by PCR using KOD plus with the primers in Table S1 and cloned into the unique *Not*I site of pENTR/D-TOPO-MpARK by In-Fusion system (Takara) to generate pENTR/D-TOPO-MpARKpro:MpARK. To introduce a silent synonymous mutation for complementation of Mp*ark-g1* mutants, the PAM sequence of gRNA1 was mutagenized by KOD mutagenesis kit (Toyobo) and primers shown in Table S1. The resulting entry vector was used in the LR reaction with the Gateway binary vector pMpGWB307^38^ to generate MpARKpro:MpARK-Citrine construct. This was introduced into regenerating thalli of the Mp*ark-g1* mutants by the method described in ref. 55. Transformants were selected on the B5 medium containing 0.5 μM chlorsulfuron and 100 µg/ml claforan.

To generate a rigor mutation (T172N), pENTR/D-TOPO-MpARKpro:MpARK with a mutation in the PAM sequence was subjected to inverse PCR using KOD mutagenesis kit (Toyobo) and primers shown in Table S1. To remove the armadillo repeats, pENTR/D-TOPO-MpARKpro:MpARK with a PAM mutation was subjected to inverse PCR using KOD mutagenesis kit (Toyobo) and primers shown in Table S1. The resulting entry vectors were subjected to the LR reaction with the Gateway binary vector pMpGWB307^38^ to generate MpARKpro:MpARK^T172N^-Citrine and MpARKpro:MpARK^dARM^-Citrine constructs. These were introduced into regenerating thalli of the Mp*ark-g1* mutants by the method described in ref. 55. Transformants were selected on the B5 medium containing 0.5 μM chlorsulfuron and 100 µg/ml claforan.

To construct MpARKpro:Citrine-NLS, a part of MpARK cDNA spanning from the 21st amino acid (Arg21) to the 883rd amino acid (Tyr883) in pENTR/D-TOPO-MpARKpro:MpARK was removed by inverse PCR using KOD mutagenesis kit (Toyobo) and primers shown in Table S1. The resulting entry vector, including a 3.4-kb region upstream of the initiation codon and 60-b coding region corresponding to the 20-amino-acid sequence of MpARK, was subjected to the LR reaction with the Gateway binary vector pMpGWB115^38^ to generate MpARKpro:Citrine-NLS construct, in which Citrine-NLS was translationally fused with the first 20-amino-acid sequence of MpARK. MpARKpro:Citrine-NLS construct was introduced into wild type F1 sporelings as previously described^37^. Transformants were selected with 10 µg/ml hygromycin B and 100 µg/ml claforan.

To construct CaMV35Spro:MpEB1a-Citrine, Mp*EB1a* genomic CDS was amplified from Tak-1 genomic DNA by PCR using KOD plus with the primers in Table S1 and cloned into pENTR/D-TOPO. This entry vector was used in the LR reaction with the Gateway binary vector pMpGWB306^38^ to generate CaMV35Spro:MpEB1a-Citrine construct, in which Citrine was translationally fused with MpEB1b. CaMV35Spro:MpEB1a-Citrine was introduced into F1 wild-type sporelings derived from crosses between Tak-1 and Tak-2 or into regenerating thalli of Mp*ark* mutants as described above. Transformants were selected with 0.5 μM chlorsulfuron and 100 µg/ml claforan.

CaMV35Spro:Citrine-MpTUB2 was constructed by LR reaction between pENTR/D-TOPO-MpTUB2^39^ and pMpGWB305^38^. CaMV35Spro:Citrine-MpTUB2 was introduced into F1 wild-type sporelings derived from crosses between Tak-1 and Tak-2 or into regenerating thalli of Mp*ark* mutants as described above. Transformants were selected with 0.5 μM chlorsulfuron and 100 µg/ml claforan.

To construct MpARKpro:MpARK-TagRFP, pENTR/D-TOPO-MpARKpro:MpARK (with a silent mutation in the PAM sequence of gRNA1) was used in the LR reaction with the Gateway binary vector pMpGWB326^38^ to generate MpARKpro:MpARK-TagRFP construct. This was introduced into regenerating thalli of CaMV35Spro:Citrine-MpTUB2^39^ (pMpGWB106-MpTUB2) or CaMV35Spro:MpEB1a-Citrine by the method described above. Transformants were selected on the B5 medium containing 0.5 μM chlorsulfuron and 100 µg/ml claforan.

To construct CaMV35Spro:MpARK-Citrine, pENTR/D-TOPO-MpARK was used in the LR reaction with the Gateway binary vector pMpGWB306^38^. This was introduced into F1 wild-type sporelings derived from crosses between Tak-1 and Tak-2 by the method described above. Transformants were selected on the B5 medium containing 0.5 μM chlorsulfuron and 100 µg/ml claforan.

To visualize nuclear movement, MpNEK1pro:Citrine-NLS was constructed by LR reaction between pENTR/D-TOPO-MpNEK1pro^39^ and pMpGWB315^38^. MpNEK1pro:Citrine-NLS construct was introduced into F1 wild-type sporelings derived from crosses between Tak-1 and Tak-2 or into regenerating thalli of Mp*ark* mutants as described above. Transformants were selected on the B5 medium containing 0.5 μM chlorsulfuron and 100 µg/ml claforan.

To visualize TGN, pMpGWB301 harboring MpSYP2pro:Citrine-MpSYP6A^41^ was introduced into F1 wild-type sporelings derived from crosses between Tak-1 and Tak-2 or into regenerating thalli of Mp*ark* mutants as described above. Transformants were selected on the B5 medium containing 0.5 μM chlorsulfuron and 100 µg/ml claforan.

To construct MpNEK1pro:MpNEK1-TagRFP, pENTR/D-TOPO-MpNEK1pro:MpNEK1^39^ was used in the LR reaction with the Gateway binary vector pMpGWB426^38^ to generate MpNEK1pro:MpNEK1-TagRFP construct. This was introduced into regenerating thalli of MpARKpro:MpARK-Citrine by the method described above. Transformants were selected on the B5 medium containing 5 µg/ml G418 and 100 µg/ml claforan.

For the localization analysis of MpNEK1, pMpGWB307-MpNEK1pro:MpNEK1^39^ was introduced into F1 wild-type sporelings derived from crosses between Tak-1 and Tak-2 or into regenerating thalli of Mp*ark* mutants as described above. Transformants were selected on the B5 medium containing 0.5 μM chlorsulfuron and 100 µg/ml claforan.

To construct AtNEK6pro:MpARK-GFP, pENTR/D-TOPO-MpARK was used in the LR reaction with the Gateway binary vector pGWB504^56^ containing *NEK6* promoter (details of this construct will be described elsewhere by H.M.) to generate AtNEK6pro:MpARK-GFP construct. This was introduced into *ark1-1 ark2-1 ark3-1* triple mutant^23^ of *A. thaliana* by floral dip method^57^. Transformants were selected on the MS medium containing 10 µg/ml hygromycin B and 100 µg/ml claforan.

### Microscopy

Plants were observed and photographed by a stereomicroscope S8 Apo (Leica Microsystems, Wetzlar, Germany) equipped with a CCD camera EC3 (Leica Microsystems) or by a single-lens reflex camera D5600 (Nikon, Tokyo, Japan). Morphology of rhizoids was observed by S8 Apo equipped with EC3 or a light microscope DM5000B (Leica) equipped with a CCD camera DFC500 (Leica). Rhizoid length was quantified by ImageJ software (NIH, USA).

For live imaging of rhizoids, the B5 agar medium was poured and solidified in a glass bottom dish (35 mm in diameter x 10 mm in thickness). The central part of the medium on the bottom slide glass was removed by tweezers and was supplied with about 200 µL of melted B5 agar medium to form a thin solidified medium. The gemmae or thalli were planted on the center of medium and grown for 2-7 days under continuous white light at 22°C. The details of this method will be described elsewhere by A.K. and H.M.

The gemmalings and thalli were observed under FV1200 confocal laser-scanning microscope (Olympus) equipped with a high-sensitivity GaAsP detector and silicone oil objective lenses (30 x, numerical aperture (NA) = 1.05; 60 x, NA = 1.3, Olympus) or FV3000 (Olympus) equipped with high-sensitivity GaAsP detectors and oil immersion objective lenses (40x, NA = 1.4; 60x, NA = 1.42, Olympus). Silicone oil (SIL300CS, Olympus) or immersion oil (F30CC, Olympus) was used as immersion medium for objective lens. The samples were excited at 473 nm and 559 nm (laser diode). The emission was separated using a FV12-MHSY SDM560 filter (490-540 nm, 575-675 nm, Olympus) in FV1200.

The samples were also observed using an LSM780 confocal microscope (Carl Zeiss) equipped with an oil immersion lens (40x, NA = 1.4) and lambda and Airyscan detectors^58^. For spectral imaging, the samples were excited at 488 nm (Argon 488) and 561 nm (DPSS 561-10), and emissions between 482 and 659 nm were collected. For high-resolution imaging using the Airyscan detector, samples were excited at 488 and 561 nm, and the emission was separated using a BP495-550 + BP570-630 filter. The images were acquired using line scanning. Spectral unmixing and Airy processing of the obtained images were performed using ZEN2.3 SP1 software (Carl Zeiss).

Obtained images were analyzed using ImageJ (National Institutes of Health, USA) and ImageJ Fiji^59^ with ImageJ plugins such as LPixel plugins (LPixel, Japan). MpARK dots and MpEB1a comets were tracked by the MTrackJ plugin to quantify their velocity. The catastrophe of microtubule plus-ends was calculated by the inverse of the duration time of MpEB1a comets according to ref. 7. Because plastids exhibited back and forth movement, their motion was analyzed by the kymograph plugin to measure the tip-directed moving distance and migration time. The velocity of plastids were calculated by dividing the tip-directed moving distance by the migration time. Because the nucleus showed constant tip-directed movement, nucleus motility was quantified by the line tool, which was applied from the center of nucleus in the first flame to that in the last flame. The velocity of the nucleus were calculated by dividing the tip-directed moving distance by the migration time. The distances from the tip-side edge of plastids and nucleus to the tip of rhizoids were quantified by the line tool. TGNs labelled with Citrine-SYP6A were tracked by the MTrackJ plugin. The velocity of TGNs were calculated by dividing the distance from the first position to the last position (values of Max D2S in MTrackJ) by the migration time to exclude the effect of Brownian motion, which was frequently observed in the TGNs of Mp*ark* mutants.

### RT-qPCR

Total RNA was isolated by NucleoSpin^®^ RNA Plant RNA isolation kit (Takara Bio, Kusatsu, Japan) according to the manufacture’s instruction. RNA was quantified with a spectrophotometer DU730 (Beckman Coulter, CA, USA). Reverse transcription was performed with 0.5 µg of total RNA of each sample by ReverTra Ace reverse transcriptase (Toyobo, Osaka, Japan) according to the manufacture protocol. Quantitative real-time PCR was performed in Thermal Cycler Dice Real Time System Lite (Takara Bio) using KOD SYBR qPCR Mix (Toyobo) and gene-specific primers shown in Table S1. Transcript levels of Mp*ACT* were used as references for normalization^60^. RT-qPCR was conducted in four biological replicates.

### Phylogenetic analysis

For phylogenetic analysis of ARK kinesins, we used a conserved motor domain at the N-terminus. Sequence information was collected in Phytozome (http://phytozome.jgi.doe.gov/pz) and 1KP database (https://db.cngb.org/onekp/). To generate a maximum-likelihood bootstrap consensus tree, the sequences were aligned by MUSCLE^61^ and phylogenetic analyses were conducted in MEGA7^62^. The evolutionary history was estimated using the Maximum Likelihood method based on the JTT matrix-based model. The bootstrap consensus tree deduced from 1000 replicates is adopted to represent the evolutionary history of the proteins analyzed. The percentages of replicate trees in which the associated sequences clustered together are shown next to the branches. The branch lengths were measured in the number of substitutions per site.

### Statistical analyses

All statistical analyses were conducted using R software (R 3.6.3, R Foundation for Statistical Computing, Vienna, Austria). Tukey’s HSD test was used for comparisons of multiple sample means. The types of statistical tests, sample sizes, and significance level (*P* value) are shown in figure legend.

### Data availability

All the data that support the findings of this study are included in this article and supplementary figures and movies. Source data are provided with this paper.

## Acknowledgements

We thank Drs. Shigeo Sugano, Keishi Osakabe, and Takayuki Kohchi for providing CRISPR/Cas9 vectors, Drs. Ryuichi Nishihama, Kimitsune Ishizaki, and Takayuki Kohchi for the early MpARK sequence, pMpGWBs and MpTUB2 constructs, Dr. Tatsuya Sakai for *Arabidopsis ark* mutant seeds, Dr. Tsuyoshi Nakagawa for pGWB504, Dr. Takehiko Kanazawa for technical support in microscopy, and Dr. Gohta Goshima for discussing and sharing unpublished results. This work was supported by Grants-in-Aid from the JSPS (JSPS KAKENHI Grant Numbers 16K07403, 19K06709, 21H00370, and 23H04708 to H.M.), Ryobi Teien Memory Foundation, and Naito Foundation to H.M.

## Author contributions

A.K. and H.M. designed the experiments. A.K. performed the experimental studies. A.K. and H.M. analyzed the data. T.U., T.T. and H.M. provided research materials, reagents and equipment. A.K. and H.M. wrote the manuscript. H.M. supervised the project. All authors discussed the results and commented on the manuscript.

---

**Supplementary Fig. 1. Structure and expression of MpARK transcripts.**

(a) Splicing variants of MpARK (from Marpolbase database).

(b) RT-qPCR analysis of MpARK. Total RNA was isolated from the 21-day-old wild type plants and subjected to RT-qPCR analysis using two primer sets shown in the upper diagram (the diagram is identical to Fig. 1b). Values are shown by box plots indicating median (middle line), 25^th^, 75^th^ percentile (box) and 5th and 95th percentile (whiskers) (*n* = 4). An asterisk indicates significant difference (Wilcoxon rank sum test, P < 0.03).

**Supplementary Fig. 2. Phenotype and genotype of MpARK mutants.**

(a) Morphology of the wild type and Mp*ark-g1* mutants (left two panels: 7-day-old plants, right two panels: 21-day-old plants). Arrowheads indicate rhizoids.

(b) Morphology of rhizoids of the wild type, Mp*ark-g1* mutants, and Mp*ark-g1* complemented with the full length MpARK-Citrine (MpARKpro:MpARK-Citrine).

**Supplementary Fig. 3. Genomic sequences flanking gRNAs.**

Genomic sequences flanking target sites of MpARK (c, gRNA1; d, gRNA2). DNA and amino acid sequences of the wild type (WT) and mutant lines. Deleted bases are shown by hyphens. Inserted or substituted sequences are colored in red. Open box: PAM sequence.

**Supplementary Fig. 4. Localization and dynamics of MpARK-Citrine.**

(a) Dynamics of MpARK-Citrine in the rhizoid apex. Mp*ark-g1* complemented with the full length MpARK-Citrine was observed in a confocal microscope. This corresponds to the right cell in Movie 1.

(b) Co-localization of MpARK-TagRFP and Citrine-MpTUB2 (microtubules). Confocal z-stack image of double labelled line (MpARK*pro:*MpARK*-TagRFP, CaMV35Spro:Citrine-*Mp*TUB2*).

(c) Movement of MpARK-TagRFP particle localized to the crossover of microtubules (Movie 3). The rightmost panel, trajectory of MpARK-TagRFP. Arrowheads, MpARK-TagRFP particles. Arrows, direction of movement.

(d) Effects of oryzalin and latrunculin B (LatB) on the localization of MpARK-Citrine. Mp*ark-g1* complemented with MpARKpro:MpARK-Citrine was treated with 10 µM oryzalin or 1 µM latrunculin B for 15 min and observed under a confocal microscope. Arrows indicate MpARK-Citrine.

**Supplementary Fig. 5. Localization of MpNEK1-Citrine in the wild type and Mp*ark-g1* mutant.**

Confocal z-stack image of MpNEK1-Citrine (Mp*NEK1pro:*Mp*NEK1-Citrine*) in the wild type and Mp*ark-g1* mutant.

**Supplementary Fig. 6. Morphology of rhizoids of CaMV35S:MpARK-Citrine.**

Confocal z-stack image of CaMV35S:MpARK-Citrine.

**Supplementary Fig. 7. Association of MpARK-Citrine with plastids and the nucleus.**

Confocal z-stack image of Mp*ark-g1* complemented with MpARKpro:MpARK-Citrine.

**Supplementary Fig. 8. Association of MpARK-Citrine with plastids.**

Single optical section of Mp*ark-g1* complemented with MpARKpro:MpARK-Citrine. Spectral imaging combined with linear unmixing was used to separate MpARK-Citrine signal and plastid autofluorescence. The lower three panels are identical to the uppermost image surrounded by a dotted rectangle. The brackets indicate association of MpARK-Citrine and plastids.

**Supplementary Fig. 9. Involvement of MpARK in organelle transport.**

(a) Localization of mitochondria stained by DiOC6 in rhizoids of the wild type and Mp*ark-g1*. The wild type and Mp*ark-g1* were stained with 1 µg/mL DiOC6 for 5 min and observed under a confocal microscope (single optical section).

(b) Quantification of fluorescent density of DiOC6-stained mitochondria at the apical regions of rhizoids in the wild type and Mp*ark-g1*. The apical region is 0-10 µm region away from the rhizoid apex. Values are shown by box plots indicating median (middle line), 25^th^, 75^th^ percentile (box) and 5th and 95th percentile (whiskers) (*n* = 6 rhizoids in the wild type and 11 rhizoids in Mp*ark-g1*). An asterisk indicates significant difference from the value of the wild type (Wilcoxon rank sum test, P < 0.03).

(c) Localization of vesicles stained by BODIPY-BFA in rhizoids of the wild type and Mp*ark-g1*. The wild type and Mp*ark-g1* were stained with 0.5 µM BODIPY-BFA for 5 min and observed under a confocal microscope (single optical section in the upper and middle panels, the lower panel is a z-stack image of the wild type showing punctate staining pattern).

(d) Quantification of fluorescent density of BODIPY-BFA-stained vesicles at the apical regions of rhizoids in the wild type and Mp*ark-g1*. The apical region is 0-10 µm region away from the rhizoid apex. Values are shown by box plots indicating median (middle line), 25^th^, 75^th^ percentile (box) and 5th and 95th percentile (whiskers) (*n* = 7 rhizoids in the wild type and 11 rhizoids in Mp*ark-g1*). An asterisk indicates significant difference from the value of the wild type (Wilcoxon rank sum test, P < 0.01).

**Supplementary Movie 1**. Dynamics of MpARK-Citrine in the apical dome of rhizoids. Time-lapse of rhizoids of Mp*ark-g1* complemented with MpARKpro:MpARK-Citrine. Green, MpARK-Citrine. Magenta, plastid autofluorescence. Images were acquired every 6 seconds (25 fps).

**Supplementary Movie 2**. Dynamics of MpARK-Citrine in a rhizoid. Time-lapse of a rhizoid of Mp*ark-g1* complemented with MpARKpro:MpARK-Citrine. Green, MpARK-Citrine. Magenta, plastid autofluorescence. Images were acquired every 5 seconds (25 fps).

**Supplementary Movie 3**. Z-series image of a rhizoid expressing CaMV35Spro:Citrine-MpTUB2 (green) and MpARKpro:MpARK-TagRFP (magenta). Images were acquired every 1 µm along the z axis (5 fps).

**Supplementary Movie 4**. Dynamics of MpARK in a rhizoid apex. Time-lapse of a rhizoid apex expressing CaMV35Spro:Citrine-MpTUB2 (green) and MpARKpro:MpARK-TagRFP (magenta). Images were acquired every 3 seconds (20 fps).

**Supplementary Movie 5**. Z-series image of a rhizoid expressing CaMV35Spro:MpEB1a-Citrine (green) and MpARKpro:MpARK-TagRFP (magenta). Images were acquired every 1 µm along the z axis (7 fps).

**Supplementary Movie 6**. Z-series image of a rhizoid expressing CaMV35Spro:MpEB1a-Citrine (green) and MpARKpro:MpARK-TagRFP (magenta). Images were acquired every 0.4 µm along the z axis (10 fps).

**Supplementary Movie 7**. Z-series image of a rhizoid expressing CaMV35Spro:Citrine-MpTUB2 (green) and MpARKpro:MpARK-TagRFP (magenta) obtained by an enhanced-resolution confocal microscope equipped with Airyscan detector. Images were acquired every 1 µm along the z axis (2 fps).

**Supplementary Movie 8**. Z-series image of a rhizoid expressing CaMV35Spro:Citrine-MpTUB2 (green) and MpARKpro:MpARK-TagRFP (magenta) obtained by an enhanced-resolution confocal microscope equipped with Airyscan detector. Images were acquired every 1 µm along the z axis (2 fps).

**Supplementary Movie 9**. Dynamics of MpARK and MpNEK1 in a rhizoid. Time-lapse of a rhizoid expressing MpARKpro:MpARK-Citrine (green) and MpNEK1pro:MpNEK1-TagRFP (magenta). Images were acquired every 3 seconds (20 fps).

**Supplementary Movie 10.** Dynamics of MpARK and MpNEK1 in a rhizoid. Time-lapse of a rhizoid expressing MpARKpro:MpARK-Citrine (green) and MpNEK1pro:MpNEK1-TagRFP (magenta). Images were acquired every 4 seconds (20 fps).

**Supplementary Movie 11.** Dynamics of MpEB1a-Citrine in the wild-type rhizoids. Time-lapse of rhizoids of the wild type with CaMV35Spro:MpEB1a-Citrine. Images were acquired every 5 seconds (20 fps).

**Supplementary Movie 12.** Dynamics of MpEB1a-Citrine in the Mp*ark-g1* rhizoid. Time-lapse of a rhizoid of the Mp*ark-g1* mutant with CaMV35Spro:MpEB1a-Citrine. Images were acquired every 4 seconds (20 fps).

**Supplementary Movie 13.** Motility of the nucleus (green) and plastids (magenta) in the wild-type rhizoid. Time-lapse of a rhizoid of the wild type with Mp*NEK1pro:Citrine-NLS*. Images were acquired every 8 seconds (20 fps).

**Supplementary Movie 14.** Motility of the nucleus (green) and plastids (magenta) in the Mp*ark-g1* mutant rhizoid. Time-lapse of a rhizoid of the Mp*ark-g1* mutant with Mp*NEK1pro:Citrine-NLS*. Images were acquired every 15 seconds (10 fps).

**Supplementary Movie 15.** Motility of TGN (green) and plastids (magenta) in the wild-type rhizoid. Time-lapse of a rhizoid of the wild type expressing Citrine-MpSYP6A. Images were acquired every 6 seconds (20 fps).

**Supplementary Movie 16.** Motility of TGN in the Mp*ark-g1* mutant rhizoid. Time-lapse of a rhizoid of the Mp*ark-g1* mutant expressing Citrine-MpSYP6A. Images were acquired every 5 seconds (20 fps).

