## Supplementary material for "Plant-specific armadillo repeat kinesin directs organelle transport and microtubule convergence to promote tip growth": Suppl Fig & Table

Supplementary Fig. 1

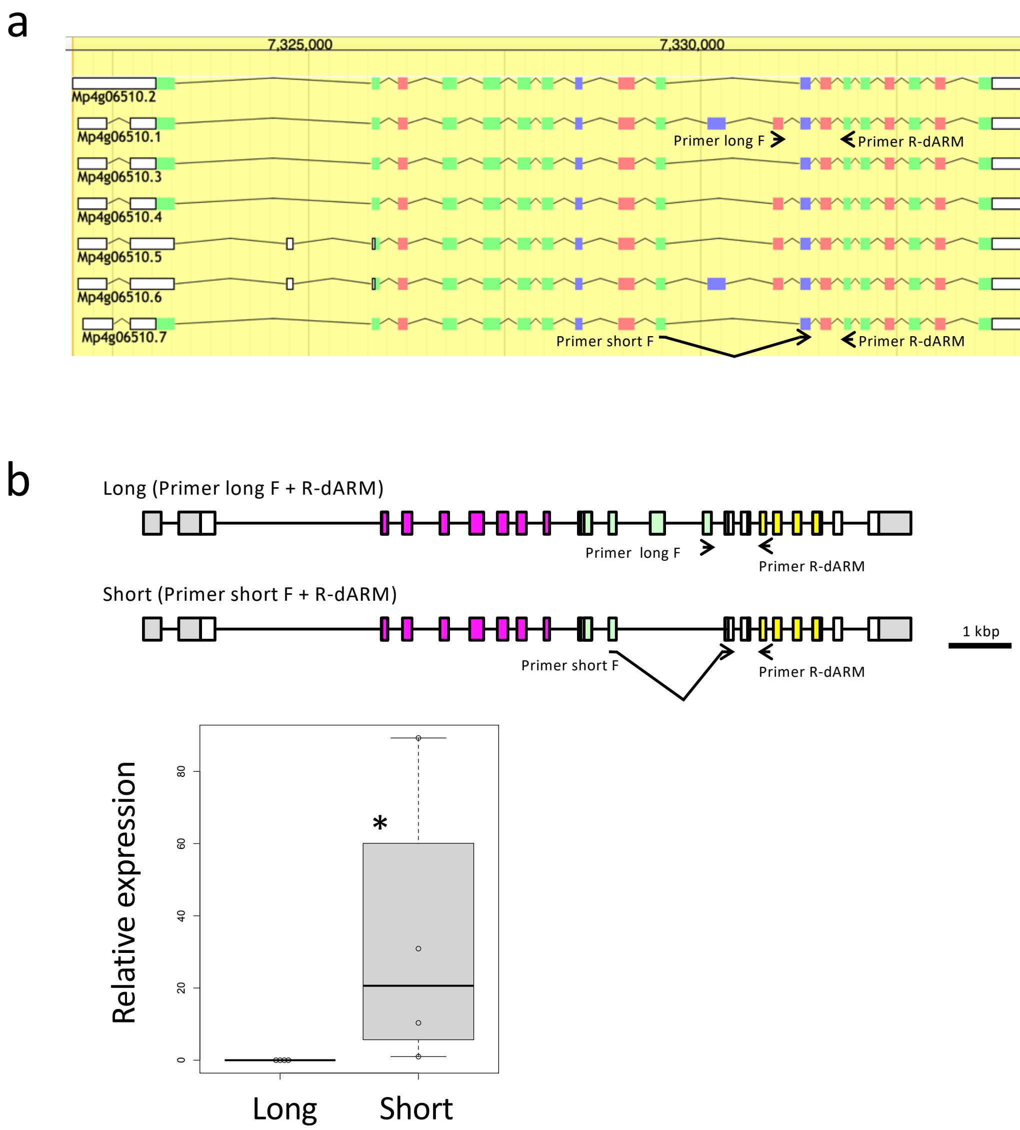


**Supplementary Fig. 1. Structure and expression of MpARK transcripts.**
(a) Splicing variants of MpARK (from Marpolbase database).
(b) RT-qPCR analysis of MpARK. Total RNA was isolated from the 21-day-old wild type plants and subjected to RT-qPCR analysis using two primer sets shown in the upper diagram (the diagram is identical to Fig. 1b). Values are shown by box plots indicating median (middle line), 25th, 75th percentile (box) and 5th and 95th percentile (whiskers) ($n$ = 4). An asterisk indicates significant difference (Wilcoxon rank sum test, $P < 0.03$).

Supplementary Fig. 2

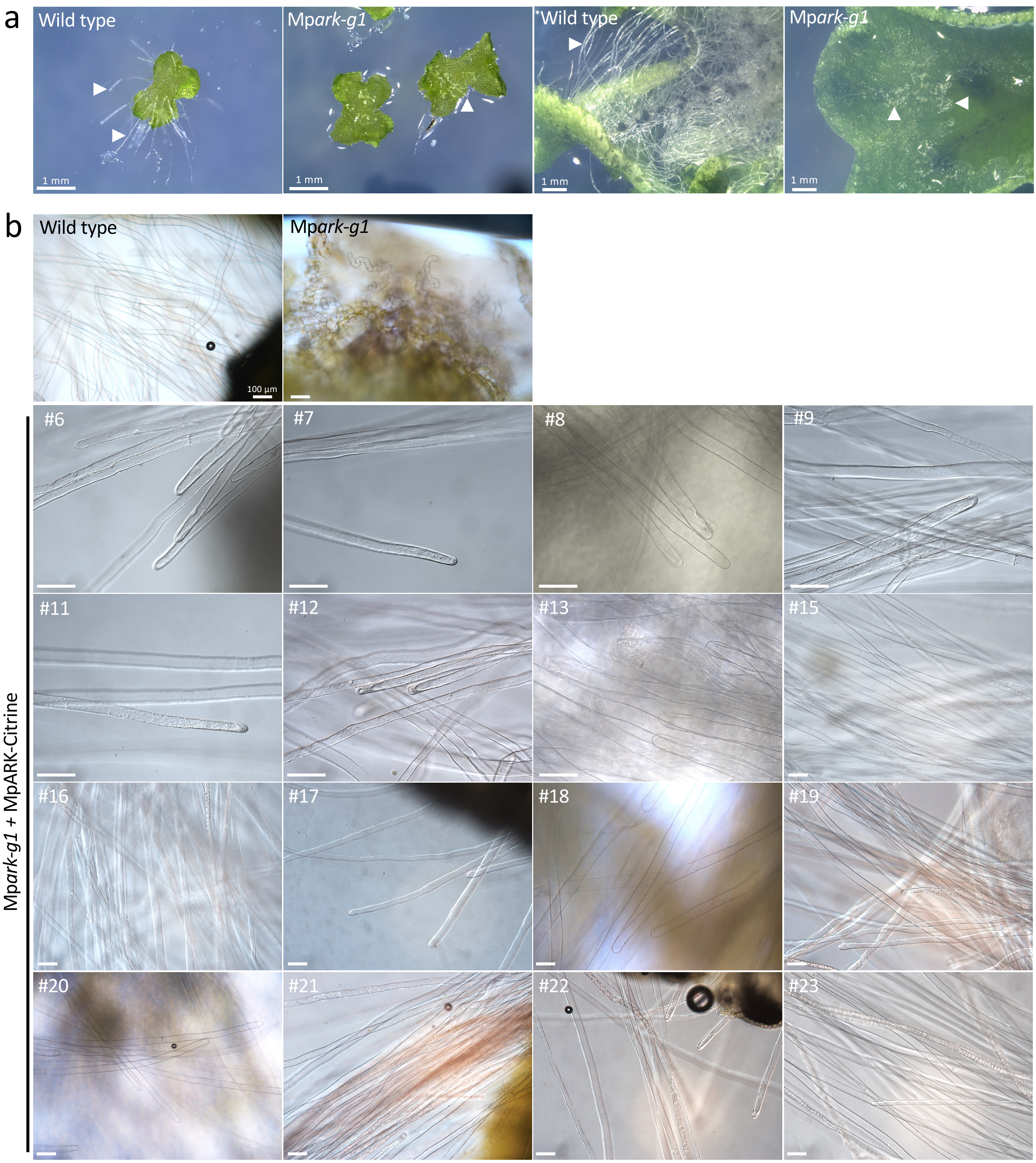


**Supplementary Fig. 2. Phenotype and genotype of MpARK mutants.**
(a) Morphology of the wild type and Mp*ark-g1* mutants (left two panels: 7-day-old plants, right two panels: 21-day-old plants). Arrowheads indicate rhizoids.
(b) Morphology of rhizoids of the wild type, Mp*ark-g1* mutants, and Mp*ark-g1* complemented with the full length MpARK-Citrine (MpARKpro:MpARK-Citrine).

Supplementary Fig. 3

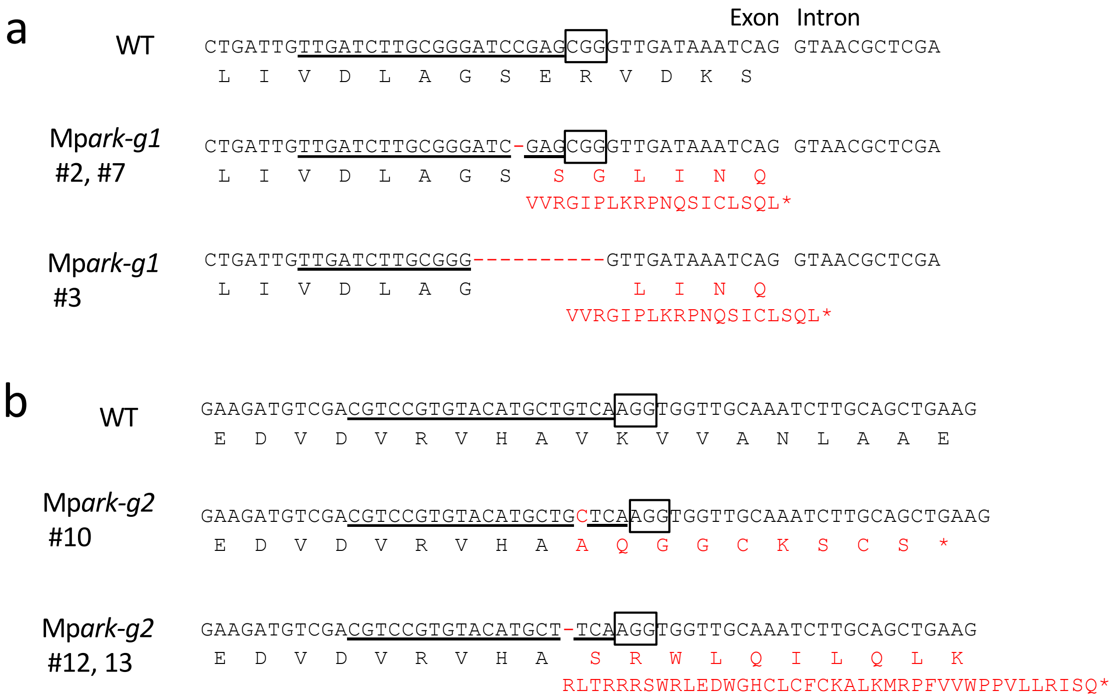


**Supplementary Fig. 3. Genomic sequences flanking gRNAs.**
Genomic sequences flanking target sites of MpARK (c, gRNA1; d, gRNA2). DNA and amino acid sequences of the wild type (WT) and mutant lines. Deleted bases are shown by hyphens. Inserted or substituted sequences are colored in red. Open box: PAM sequence.

Supplementary Fig. 4

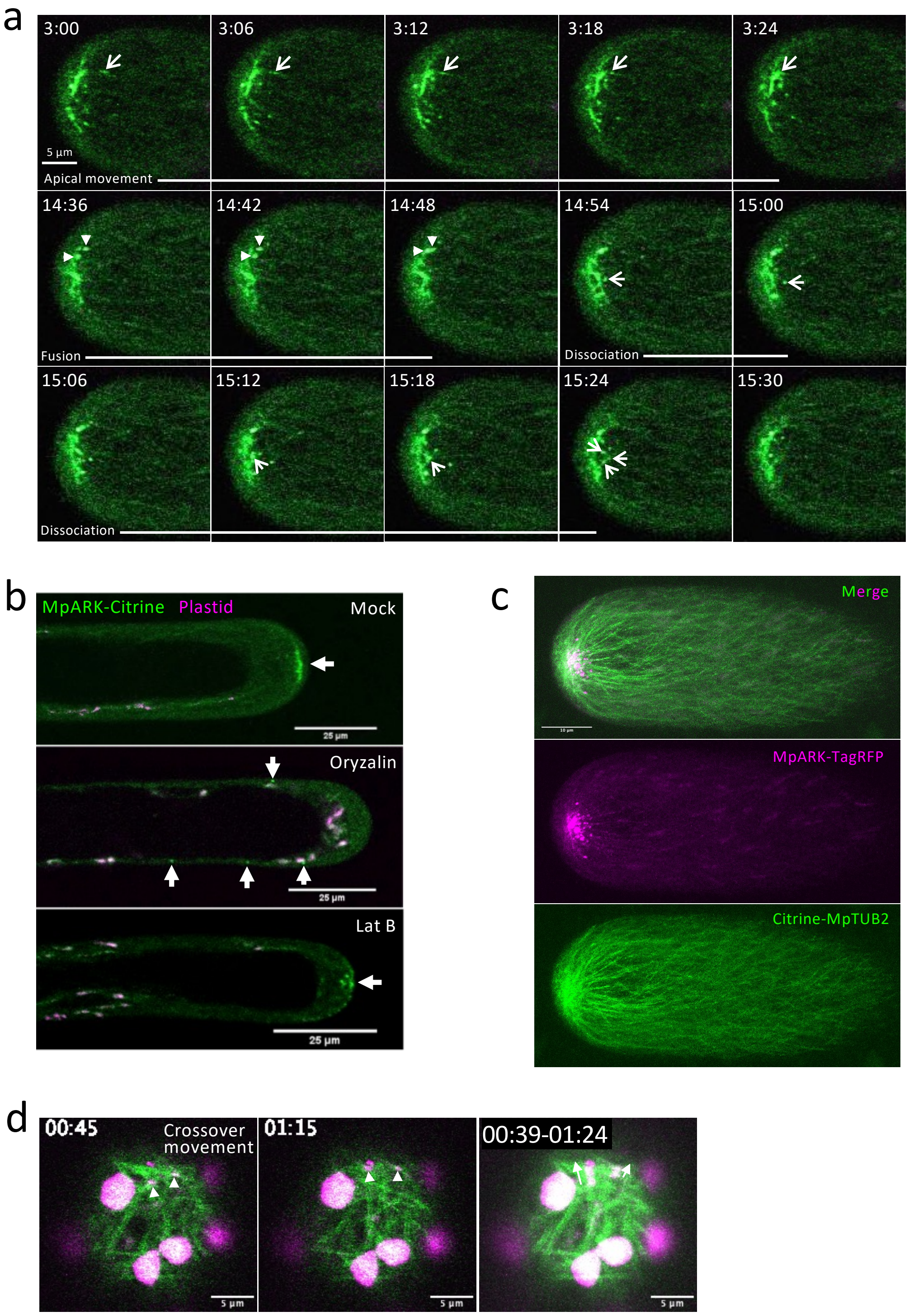


**Supplementary Fig. 4. Localization and dynamics of MpARK-Citrine.**
(a) Dynamics of MpARK-Citrine in the rhizoid apex. Mp*ark-g1* complemented with the full length MpARK-Citrine was observed in a confocal microscope. This corresponds to the right cell in Movie 1.
(b) Co-localization of MpARK-TagRFP and Citrine-MpTUB2 (microtubules). Confocal z-stack image of double labelled line (MpARK*pro:*MpARK-*TagRFP, CaMV35Spro:Citrine*-Mp*TUB2*).
(c) Movement of MpARK-TagRFP particle localized to the crossover of microtubules (Movie 3). The rightmost panel, trajectory of MpARK-TagRFP. Arrowheads, MpARK-TagRFP particles. Arrows, direction of movement.
(d) Effects of oryzalin and latrunculin B (LatB) on the localization of MpARK-Citrine. Mp*ark-g1* complemented with MpARKpro:MpARK-Citrine was treated with 10 μM oryzalin or 1 μM latrunculin B for 15 min and observed under a confocal microscope. Arrows indicate MpARK-Citrine.

Supplementary Fig. 5

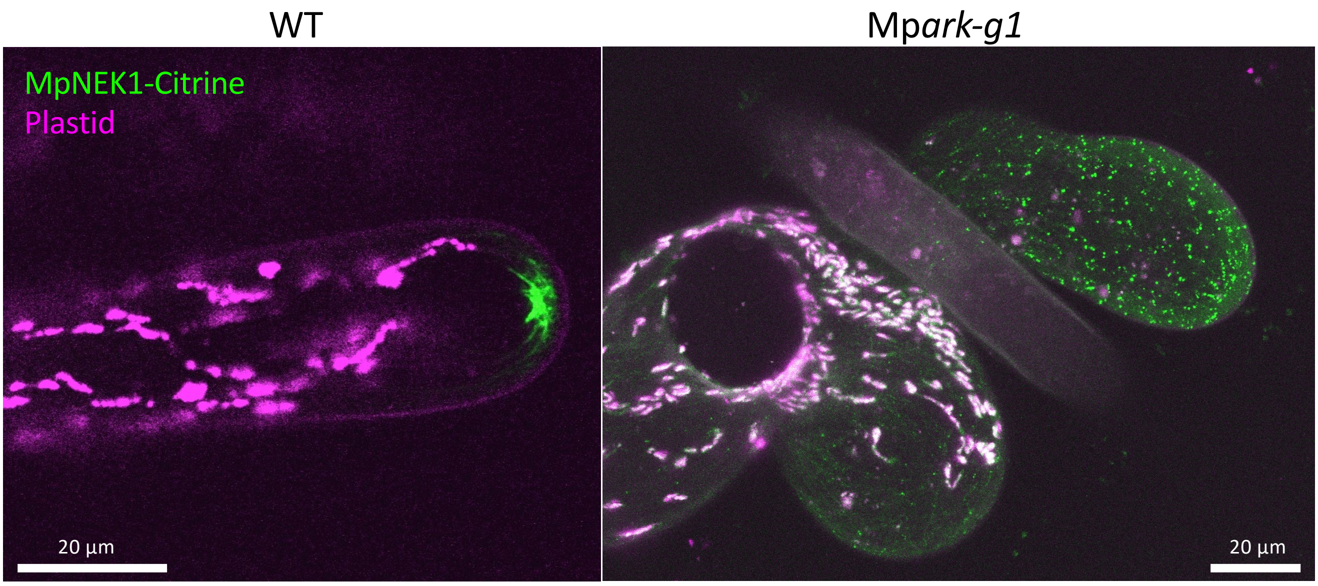


**Supplementary Fig. 5. Localization of MpNEK1-Citrine in the wild type and Mp*ark-g1* mutant.**
Confocal z-stack image of MpNEK1-Citrine (Mp*NEK1pro*:Mp*NEK1-Citrine*) in the wild type and Mp*ark-g1* mutant.

Supplementary Fig. 6

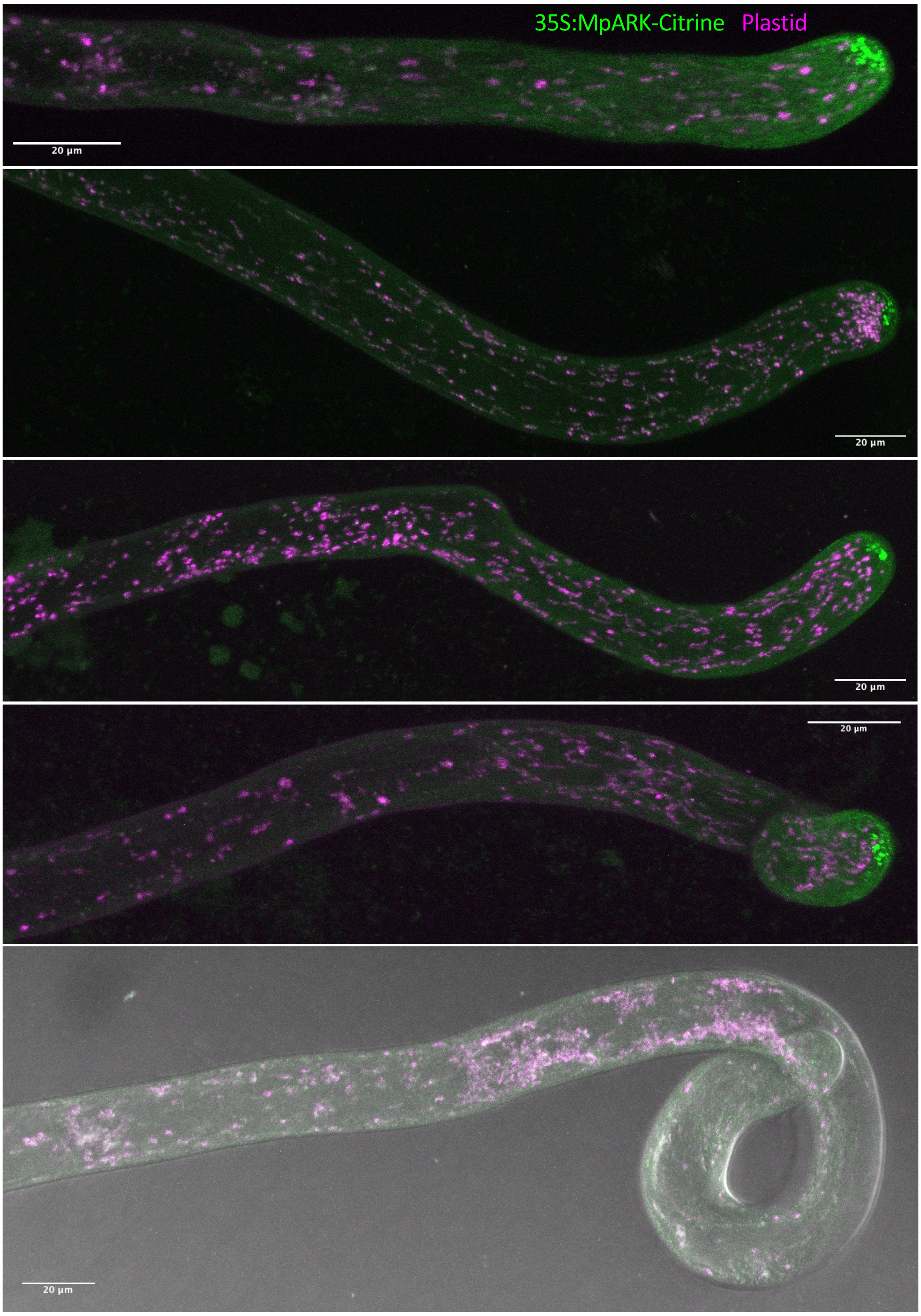


**Supplementary Fig. 6. Morphology of rhizoids of CaMV35S:MpARK-Citrine.**
Confocal z-stack image of CaMV35S:MpARK-Citrine.

Supplementary Fig. 7

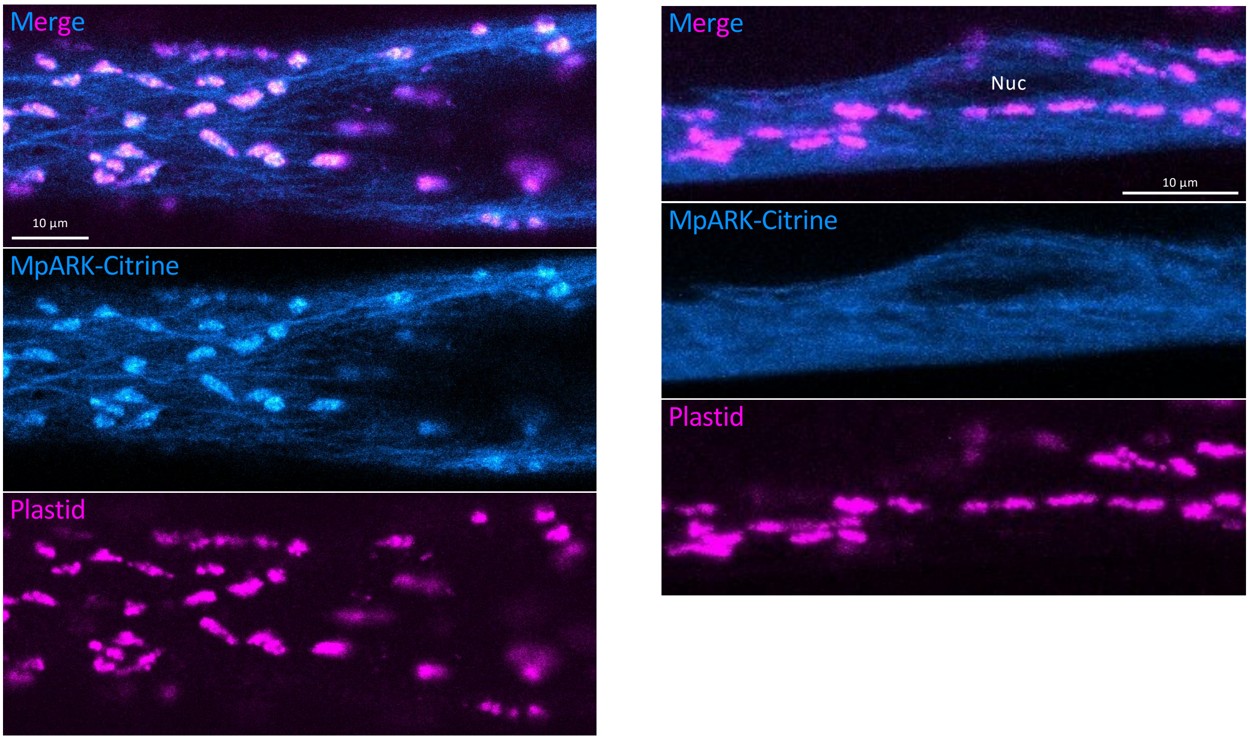


**Supplementary Fig. 7. Association of MpARK-Citrine with plastids and the nucleus.**
Confocal z-stack image of Mp*ark-g1* complemented with MpARKpro:MpARK-Citrine.

Supplementary Fig. 8

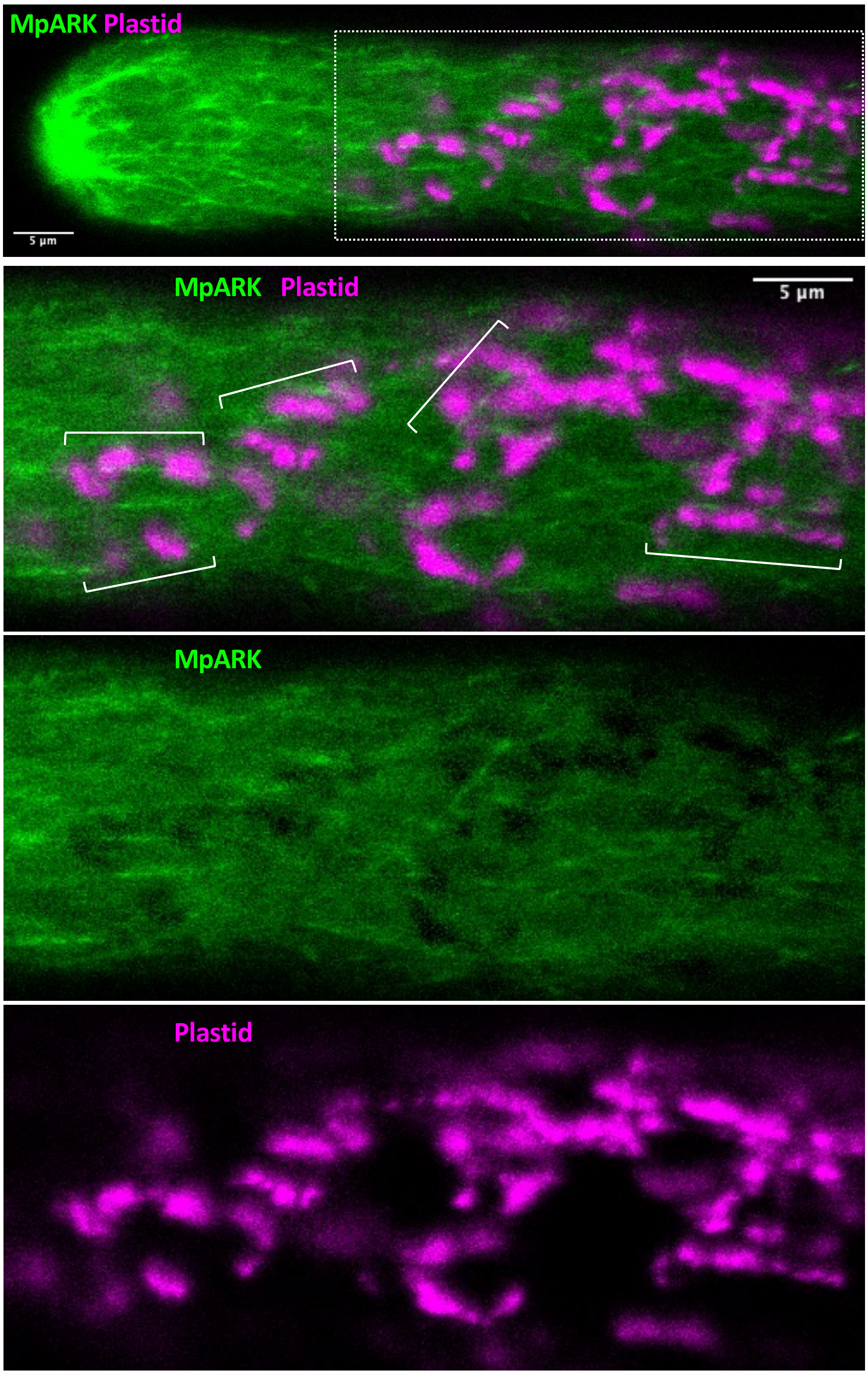


**Supplementary Fig. 8. Association of MpARK-Citrine with plastids.**
Single optical section of Mp*ark-g1* complemented with MpARKpro:MpARK-Citrine. Spectral imaging combined with linear unmixing was used to separate MpARK-Citrine signal and plastid autofluorescence. The lower three panels are identical to the uppermost image surrounded by a dotted rectangle. The brackets indicate association of MpARK-Citrine and plastids.

Supplementary Fig. 9

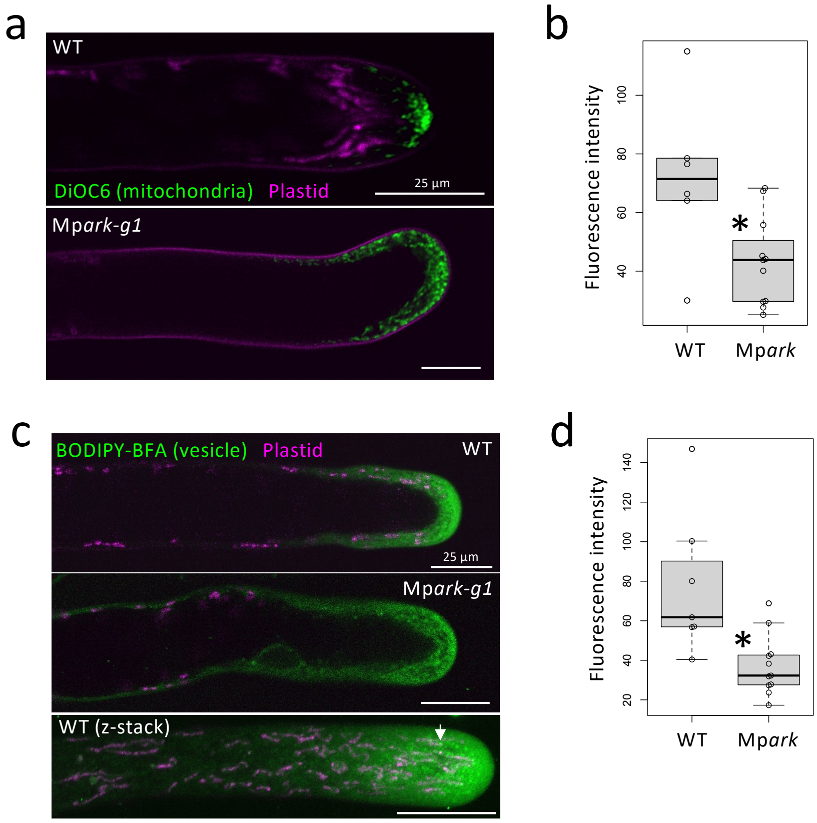


**Supplementary Fig. 9. Involvement of MpARK in organelle transport.**

(a) Localization of mitochondria stained by DiOC6 in rhizoids of the wild type and Mp*ark-g1*. The wild type and Mp*ark-g1* were stained with 1 µg/mL DiOC6 for 5 min and observed under a confocal microscope (single optical section).

(b) Quantification of fluorescent density of DiOC6-stained mitochondria at the apical regions of rhizoids in the wild type and Mp*ark-g1*. The apical region is 0-10 µm region away from the rhizoid apex. Values are shown by box plots indicating median (middle line), 25th, 75th percentile (box) and 5th and 95th percentile (whiskers) ($n$ = 6 rhizoids in the wild type and 11 rhizoids in Mp*ark-g1*). An asterisk indicates significant difference from the value of the wild type (Wilcoxon rank sum test, $P < 0.03$).

(c) Localization of vesicles stained by BODIPY-BFA in rhizoids of the wild type and Mp*ark-g1*. The wild type and Mp*ark-g1* were stained with 0.5 µM BODIPY-BFA for 5 min and observed under a confocal microscope (single optical section in the upper and middle panels, the lower panel is a z-stack image of the wild type showing punctate staining pattern).

(d) Quantification of fluorescent density of BODIPY-BFA-stained vesicles at the apical regions of rhizoids in the wild type and Mp*ark-g1*. The apical region is 0-10 µm region away from the rhizoid apex. Values are shown by box plots indicating median (middle line), 25th, 75th percentile (box) and 5th and 95th percentile (whiskers) ($n$ = 7 rhizoids in the wild type and 11 rhizoids in Mp*ark-g1*). An asterisk indicates significant difference from the value of the wild type (Wilcoxon rank sum test, $P < 0.01$).

Table S1. Primers used in this study

| Name | Sequence | Purpose |
|---|---|---|
| MpARK1-F-gRNA1 | CTCGTTGATCTTGCGGGATCCGAG | Mutagenesis of motor domain |
| MpARK1-R-gRNA1 | AAACCTCGGATCCCGCAAGATCAA | Mutagenesis of motor domain |
| MpARK1-F-gRNA2 | ctcgCGTCCGTGTACATGCTGTCA | Mutagenesis of armadillo repeat domain |
| MpARK1-R-gRNA2 | aaacTGACAGCATGTACACGGACG | Mutagenesis of armadillo repeat domain |
| MpARK1-F2-725 | CAGTACCTAGCGCTACACAT | PCR and sequence of Mp*ark1-g1* , sequence of cDNA |
| MpARK1-R2-1271 | GCTCTTTGACCGAACATGAT | PCR and sequence of Mp*ark1-g1* , sequence of cDNA |
| MpARK1-F3 | GTTTCAGCAGGATCTTGCAGAAGC | Sequence of cDNA |
| MpARK1-R3 | ATGTGGACTGTTCTCAGCGAGG | PCR and sequence of Mp*ark1-g1* , sequence of cDNA |
| MpARK1-F4 | GAACTTTGAGAGAGCAGCTGACGGT | PCR and sequence of Mp*ark1-g2* , sequence of cDNA |
| MpARK1-R4 | GCCATTATCAGCTCCTGATTGGTCT | PCR and sequence of Mp*ark1-g2* , sequence of cDNA |
| MpARK1-Fstart | caccATGGCATTTGCGTCGTCCCCCCATAA | Cloning of MpARK1 cDNA |
| MpARK1-nonstop-R | GTAGACCAACTGTAACCTTTTCAATTCT | Cloning of MpARK1 cDNA |
| MpARK1pro-F-InFus | caggctccgcggccgAGGAAGCATCAGAAACTAGGAGGA | InFusion cloning of MpARK1 promoter |
| MpARK1pro-R-InFus | gtgaagggggcggccCTCGTTACCGCATTCAAATTGCC | InFusion cloning of MpARK1 promoter |
| MpARK1-g2mut-F | CGtGTTGATAAATCAGGTAGTGAGGGGC | Mutagenesis of gRNA1 PAM sequence (CGG to CGT) |
| MpARK1-g2-R | CTCGGATCCCGCAAGATCAACAAT | Mutagenesis of gRNA1 PAM sequence (CGG to CGT) |
| MpARK1-R-Gly20 | TCCTGTCGAGCGAGCGTTCAG | MpARK1pro construct by iPCR |
| MpARK1-F-T172N | aatTTCACTCTGGGACGACTTGGAGAGG | Rigor mutation |
| MpARK1-R-K171 | CTTCCCGGTACCAGTTTGACCATAA | Rigor mutation |
| MpARK1-R-dARM | TTGACCATTTATTGTATCCCTCATATG | Deletion of armadillo repeats by iPCR, RT-qPCR |
| pENTR-F1 | AAGGGTGGGCGCGCCGACCCA | Foward primer for iPCR |
| MpARK-ShortRT-F1 | TCAGCATGAGAAGAAGGAGGCAAAGC | RT-qPCR |
| MpARK-LongRT-F1 | AATCACAGCCACCCAAGCTGCCA | RT-qPCR |
| MpEB1a-F-start | caccATGGCGGCTACCAACATTGGGATGA | Cloning of MpEB1a |
| MpEB1a-R-nonstop | TACCTCCCTCATTGGAGAGCATTC | Cloning of MpEB1a |
